# From sailing to steam trawling: the evolution of bottom trawl effort in the North Sea

**DOI:** 10.64898/2026.08.24.746664

**Authors:** Adriaan D. Rijnsdorp, Floris P. Bennema, Frans Veenstra, Ole R. Eigaard, Jasmin Ann-Christine Thomassen, Ciarán McLaverty

## Abstract

Bottom trawls have been used for centuries, yet studies of their impact on marine ecosystems have largely been restricted to recent decades. Here, we reconstruct fishing effort for the international trawler fleets in the North Sea from the age of sail to early steam trawling, by synthesising historical data describing vessel numbers and specifications, gear dimensions, fishing grounds, and operational characteristics. The trawler fleet increased from ca 800 sailing vessels in the 1820s to ca 3500 at the peak in sail trawling in the 1880s. Subsequently, steam trawling fleets emerged, increasing to almost 2000 vessels in the 1910s, while sailing fleets declined. Trawling grounds, covering ca 7% of the North Sea in 1820s, expanded from coastal to offshore grounds, reaching ca 25% in the 1880s, and 46% in the 1910s after the transition to steam trawling. Using a hydro- and aerodynamic approach to model the wind conditions required for sail trawling, we show that about 55% – 80% of the time at sea was suitable for trawling, providing a new quantitative basis for estimating historical fishing effort. The surface area swept by the trawl per year increased from 35,000 km2 in 1820s to 225,000 km2 in 1880s and 500,000 km2 in 1910s, corresponding to ca 60% of present levels. The trawling intensity (swept area ratio) varied between 0.9-1.7 year^-1^ in the era of sail, increasing to ca 2.0 in the 1910s. The trawling footprint (unique area trawled) increased to 160,000 km^2^, about half the present level.

## 1. Introduction

Bottom trawling provides about one quarter of global capture-fisheries landings and is an important source of protein (Hilborn et al., 2023). As bottom trawls generally have a poor size and species selectivity and are towed across the seabed, there is concern about adverse side effects (Jennings and Kaiser, 1998; McConnaughey et al., 2020), such as the bycatch of undersized fish or unwanted organisms (Kelleher, 2005; Uhlmann et al., 2014) and the disturbance of the seabed that affect benthic organisms (Hiddink et al., 2017) and bio-geochemical processes (Tiano et al., 2022; Hiddink et al., 2023).

Studies on the impact of bottom trawling on exploited species and seabed habitats are generally restricted to recent time periods for which comprehensive, standardized, and digitized time series information on landings, fishing effort and its spatial distribution are available (Worm et al., 2009; Amoroso et al., 2018; Pitcher et al., 2022; Eigaard et al. 2017). Bottom trawling, however, has a long history (McLaverty et al., 2026). Fishers have been using dredges, beach seines or small bottom trawls to catch fish and invertebrates along the coast for centuries (Schnakenbeck, 1927; Le Danois and Beauge, 1937). A major technical problem, when targeting bottom-dwelling species with a towed net, is maintaining the horizontal net opening. As early as the 14^th^ century, there are records that the horizontal net opening was fixed with a wooden beam (Graham, 1956; de Groot, 1984; Jones, 2018a). Although the gear has been controversial since its introduction, due to its alleged impact on spawning and the food sources of the exploited species, it has continued to be used (de Groot, 1984; Thurstan et al., 2014).

In the North Sea, fishers in England and the Netherlands employed beam trawls to catch demersal fish species in coastal waters as early as the 14^th^ century (Beaujon, 1884; Schnakenbeck, 1927; Robinson, 1996; Jones, 2018a). It gained importance when fishers from Belgium and Germany started using the beam trawl in the early 1800s (Schnakenbeck, 1927; Lescrauwaet et al., 2013). The beam trawl fishery, however, was still small relative to the large drift net fisheries for herring and the line fisheries for cod and other gadoids (Poulsen, 2008; Holm et al., 2022). Most of the beam trawl catch was consumed locally and only part of the catch was processed and traded to inland cities in England, Belgium and Germany (van Neer and Pieters, 1997; Pieters et al., 2013; Bennema and Rijnsdorp, 2015). Prime fish, such as turbot, that could be kept alive in wells, were collected by traders at sea and sold in major population centers such as London (Schnakenbeck, 1927; de Vink, 2012).

With the establishment of railway connections between the fishing ports and major cities in the 19^th^ century, and supported by a range of technological innovations, the North Sea sail trawling fleets grew in numbers and expanded their fishing grounds to deeper waters at greater distance from the home ports (Alward, 1911; Robinson, 1996; Jones, 2018b). At the end of the 19^th^ century, bottom trawling effort increased further with the introduction of steam engines for propulsion. In contrast to sail trawlers, which were constrained by wind conditions and by periods when wind and tide were in opposite directions (Holdsworth, 1874), the transition to steam trawling increased fishing power by enabling more fishing days per year, longer daily trawling times, and higher towing speeds. A further technological development that significantly increased fishing power was the introduction of otter boards in the 1890s, which enabled much larger horizontal and vertical net opening (Garstang, 1900; Engelhard, 2008; Kerby et al., 2012). Despite the long history and substantial technological development of bottom trawling, quantitative estimates of historical trawling effort, and its spatial extent, remain particularly limited.

The objective of this study is to reconstruct the time series in bottom trawling effort (swept area [SA]) and intensity (swept area ratio [SAR]), and the trawling footprint (unique area trawled)of sail and early steam trawlers in the North Sea in the 19^th^ century and early 20^th^ century. We reviewed historical fisheries books and accounts from across the North Sea, and collated and digitized annual data on national vessel numbers and characteristics to estimate the development in fleet capacity. To estimate the fleet activity, we combined hydro- and aerodynamics with wind and current data to model the annual number of days with wind conditions favourable for sail-trawling. These estimates were then combined with historic information of gear dimensions and trawling speeds to estimate fleet effort and the total annual area of seabed swept by bottom trawls. To assess the trawling footprint and intensity of bottom trawling, we identified, digitized, and combined the location and extent of key 19^th^ century fishing grounds, and calculated the ratio of the area swept by bottom trawls with the area of the identified fishing grounds (SAR). Together, these results provide a historical baseline of North Sea fishing effort and a quantitative basis for assessing the impact of bottom trawling on demersal fish stocks and the seabed from a historic perspective.

## 2. Material and methods

### 2.1. Study area characteristics

The North Sea is a generally shallow sea located between 51°N and 61°N with a depth less than 200m and a depression along the coast of Norway (Norwegian Trench) with depths down to 1000 m or more (Figure 1a). The sea floor is composed of soft sediments (muddy to coarse sand) with local deposits of coarse sediments west of the Doggerbank, parallel to the continental coast of Germany and Denmark, and along the Norwegian Trench. These coarse-sediment areas correspond to the untrawlable or rough grounds shown in the early maps of the seabed (Olsen, 1878; Darmer, 1894; Close, 1920)

**Figure 1.**
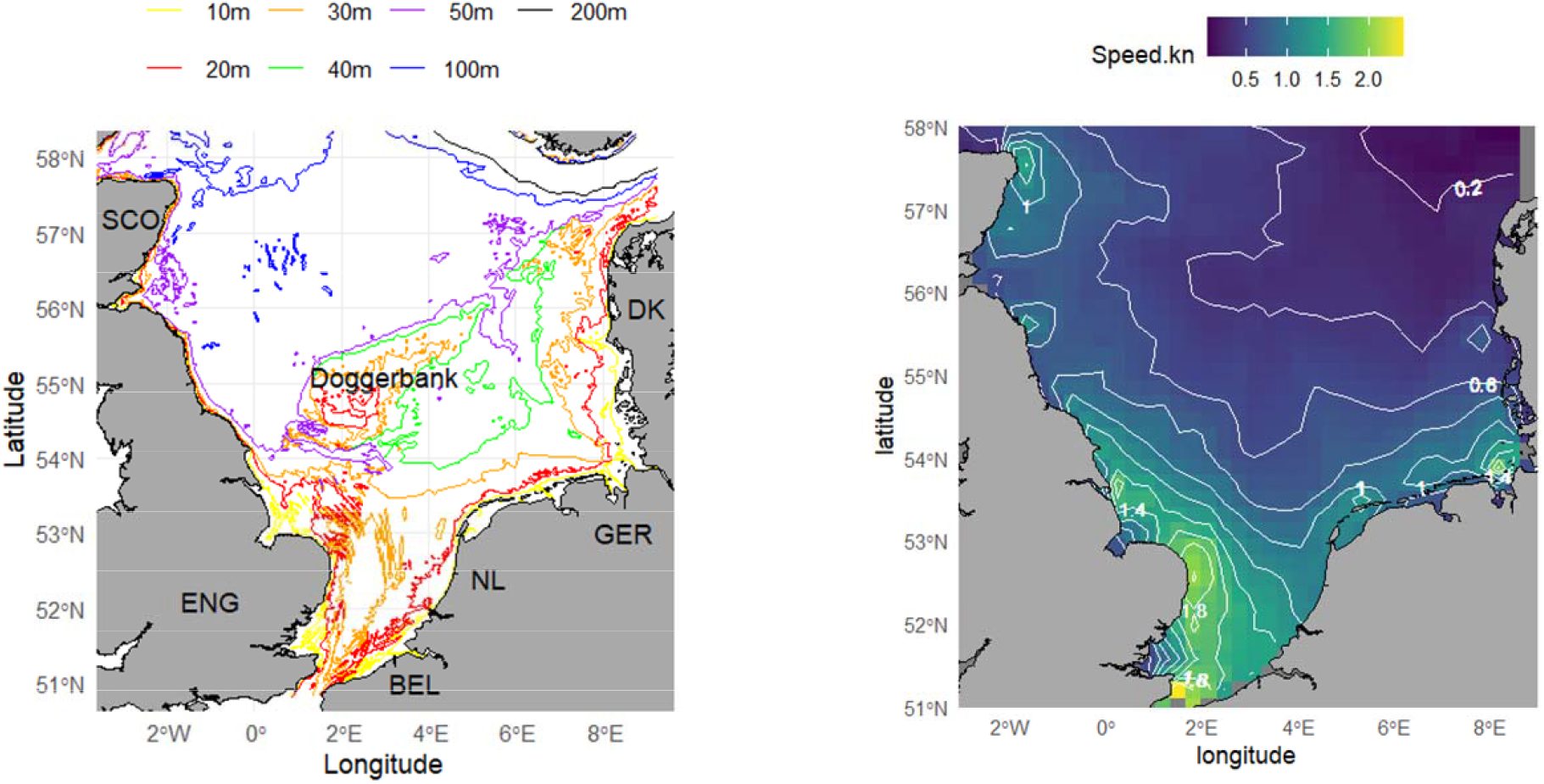
a) North Sea depth contours (DK = Denmark, GER = Germany, NL = Netherlands, BEL = Belgium, ENG = England, SCO = Scotland); b) Mean tidal speed (knot) of surface waters.s

Tidal speed ranged between 0.5 – 2.0 knot with highest speeds along the coast and decreasing going northwards (Figure 1b). Tidal speed was estimated as the mean of the hourly estimates on the u- and v-component of the surface sea water velocity in the period 01-11-2020 to 31-12-2024 for a selection of locations between 51° – 58° N and 4°W – 9°E extracted from the Copernicus database GLOBAL_ANALYSISFORECAST_PHY_001_024 (accessed on 04-06-2025). For each location, the upper 66% of values (i.e., the periods with the highest tidal speed) were extracted.

Winds over the North Sea show a dominance of westerlies in a clear seasonal pattern with lower wind force in summer (Figure 2a). Wind data was extracted from the Copernicus data base (ERA5), which provide the modelled u10- and v10-component of the wind speed (Copernicus Climate Change Service, 2023; Hersbach et al., 2023) for the period 1940 – 1979 at 2-hour intervals for a grid of locations between 51° – 60°N and 2°W – 9°E excluding the locations on land.

**Figure 2.**
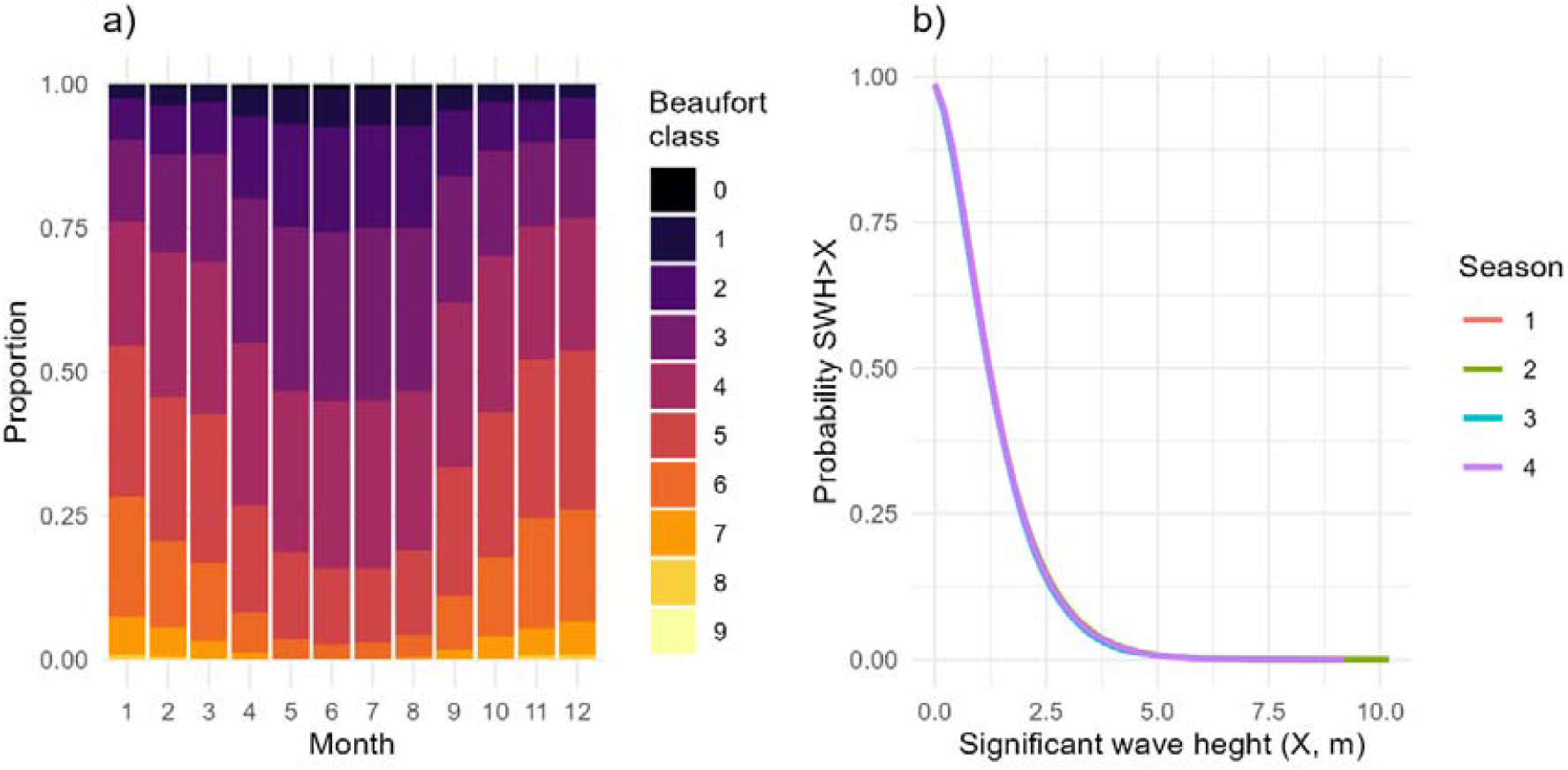
a) Probability of occurrence of wind force (Bft) by month; b) Probability that the significant wave height (SWH, m) exceeds a threshold value X.

Sea state is described as the significant wave height (SWH), which is the mean wave height (trough to crest) of the highest 33% of the waves due to winds and swell. The probability that SWH exceeds a certain value is shown in Figure 2b. Data was obtained from Copernicus data base (NWSHELF_REANALYSIS_WAV_004_015, dataset Id: MetO-NWS-WAV-RAN_202007, accessed 2025-05-30T08:53:40.707Z) for a grid of locations between 51° – 60°N and 2°W – 9°E.

### 2.2. Fishing gear

The beam trawl was the dominant fishing gear used in bottom trawling in the 19^th^ century (Schnakenbeck, 1927; March, 1953; Lescrauwaet et al., 2013). Only in Denmark, where offshore bottom trawling started relatively late (1880s), fishers used the Danish (anchor) seine (Mortensen and Strubberg, 1931). In contrast to the beam trawl used in the 20^th^ century, earlier beam trawls did not employ tickler chains to chase flatfish from the seafloor, whereas in the aft part of the bag shaped net pockets prevented fish from swimming forward and escape from the net (Collins, 1889; Hunt et al., 2024) (Figure 3). The ground rope was made of manilla and weighted with lead or chain links to prevent the rope from losing contact with the ground.

**Figure 3.**
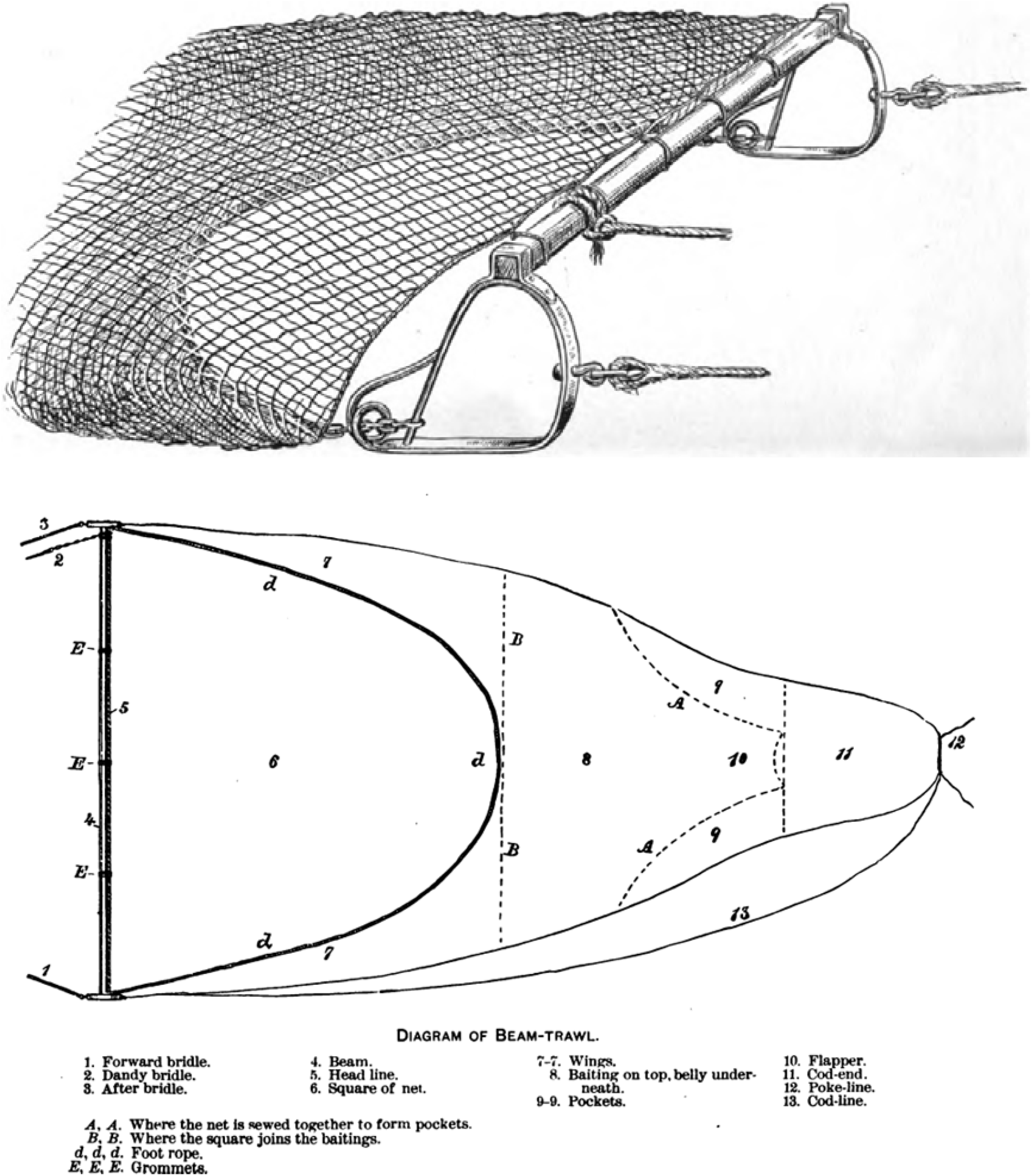
Beam trawl as used by a Grimsby smack Willy and Ada (from Collins, 1889)

Beam trawl dimensions are summarized in Table 1. Gear parameters (*Y*) were expressed as coefficients taking account of the isometric scaling relationships with vessel size (*X*) according *Y = aX*^*b*^ with b ∈ (1,2,3). Beam trawl length (BTL) was quite variable (CV=19.6%). BTL scaled with vessel length but did not differ between vessel types (Figure 4b). The resistance of a beam trawl scaled with *BTL* according to *2*.*857 BTL*^l.936^ .

**Table 1.** Beam trawl dimensions: mean, standard deviation and coefficient of variation of the coefficient (*a*) of the isometric relationship *Y = aX*^*b*^ with b ∈ (1,2,3) of various gear components.

| Table 1. Beam trawl dimensions: mean, standard deviation and coefficient of variation of the coefficient ($a$) of the isometric relationship $Y = aX^b$ with $b \in (1,2,3)$ of various gear components. | | | | | |
| --- | --- | --- | --- | --- | --- |
| Parameter<br>(Y) | Predictor<br>(X) | $b$ | Mean<br>$a$ | sd | %CV |
| Beam trawl length<br>(BTL) | LOA | 1 | 0.6054 | 0.1186 | 19.6 |
| Beam trawl length<br>diameter | BTL | 1 | 0.0158 | 0.00233 | 14.8 |
| Shoe weight | BTL | 3 | 0.0297 |  |  |
| Shoe frontal area | BTL | 2 | 0.000633 |  |  |
| Shoe sole area | BTL | 2 | 0.000449 |  |  |
| Shoe height | BTL | 1 | 0.0871 |  |  |
| Shoe width | BTL | 1 | 0.00714 |  |  |
| Rope length | BTL | 1 | 1.665 | 0.662 | 39.7 |
| Rope diameter | BTL | 1 | 0.0121 |  |  |
| Rope specific weight | Rope diameter (10cm) | 2 | 7.266 |  |  |
| Warp ratio | Depth fishing ground | 1 | 4 |  |  |
| Warp diameter | BTL | 1 | 0.011 |  |  |
| Penetration depth | constant | 1 | 0.01 |  |  |
| Net surface area | BTL | 2 | 0.151 | 0.037 | 24.5 |

**Figure 4.**
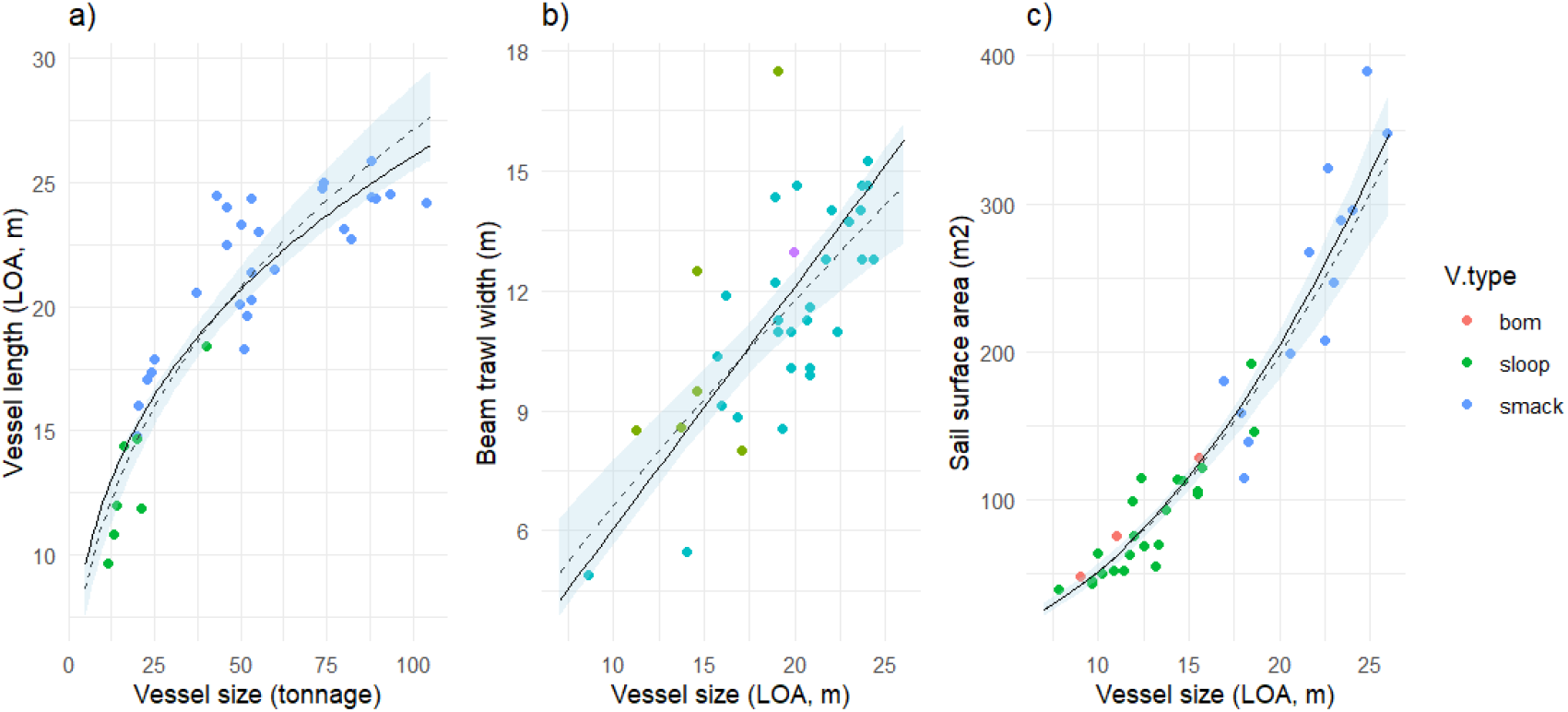
Scaling relationships sailing vessels: a) vessel length overall (LOA) - tonnage; b) beam trawl width – LOA; c) sail surface area - LOA. The dashed line and light-blue polygon show the fitted log-log relationship and 95% confidence interval. The full line shows the isometric scaling relationship (see text). V.type denotes the three main vessel types.

Steam trawlers continued fishing with a beam trawl (Hunt et al., 2024). When more powerful vessels entered the fishery in the 1890s, vessels started to use the otter trawl (Kenchington, 2018). Although there are reports that sail vessels attempted to use an otter trawl gear, it is unlikely that they could generate the thrust force required to obtain sufficient door spread (Schnakenbeck, 1927; Timmermann, 1962; de Veen, 2024).

### 2.3. Bottom trawling fleets

#### 2.3.1. Vessel types

Sail trawlers were classified according to country, vessel type, and the location from which the vessels operated, combined with information on the mode of operation and fishing grounds. Steam trawlers were classified solely by countries. Table 2 summarises the hull dimensions used to quantify the mechanics of sail trawling. Data were digitized from size-referenced drawings or taken from the text.

**Table 2.** Dimensions of sailing vessels: intercept (*a*) and slope (*b*) of the isometric relationship *Y = aX*^*b*^ of vessel parameters of three types of sailing vessels: LOA = length overall; LWL = length at the water line; Beam = vessel width; Draft = mean depth between waterline and keel; Displacement = volume of the underwater part of the hull; CM = midship coefficient; CWP = water line coefficient; CB = block coefficient; CP = prismatic coefficient; Frontal area = projected wetted surface area of the side of the hull; wetted area = surface area of the wetted hull; sail surface = surface area of the sails; tonnage = vessel size.

|  |  |  | Bomschuit |  |  | Sloop |  |  | Smack |  |  |
| --- | --- | --- | --- | --- | --- | --- | --- | --- | --- | --- | --- |
| Parameters (Y) | Predictor (X) | $b$ | Mean $a$ | sd | n | Mean $a$ | sd | n | Mean $a$ | sd | n |
| LOA | tonnage | 1/3 |  |  |  | 4.958 | 0.571 | 7 | 5.800 | 0.499 | 27 |
| LWL | LOA | 1 | 0.929 | 0.024 | 2 | 0.914 | 0.073 | 21 | 0.911 | 0.020 | 14 |
| Beam | LOA | 1 | 0.504 | 0.044 | 5 | 0.333 | 0.044 | 26 | 0.260 | 0.028 | 35 |
| Draft | LOA | 1 | 0.111 | 0.029 | 4 | 0.073 | 0.028 | 22 | 0.101 | 0.011 | 6 |
| Displacement | LWL | 3 | 0.038 | 0.006 | 3 | 0.015 | 0.006 | 5 | 0.009 | 0.004 | 12 |
| CM |  |  | 0.791 | 0.065 | 3 | 0.803 | 0.075 | 5 | 0.653 | 0.105 | 12 |
| CWP |  |  | 0.924 | 0.055 | 3 | 0.646 | 0.140 | 5 | 0.701 | 0.047 | 12 |
| CB |  |  | 0.713 | 0.048 | 3 | 0.459 | 0.120 | 5 | 0.312 | 0.115 | 12 |
| CP |  |  | 0.903 | 0.054 | 3 | 0.575 | 0.150 | 5 | 0.473 | 0.137 | 12 |
| Frontal area | LWL | 2 | 0.100 | 0.010 | 3 | 0.087 | 0.030 | 5 | 0.114 | 0.012 | 12 |
| Wetted area | LWL | 2 | 0.536 | 0.040 | 3 | 0.335 | 0.065 | 5 | 0.258 | 0.060 | 12 |
| Sail surface area | LOA | 2 | 0.582 | 0.049 | 3 | 0.501 | 0.105 | 22 | 0.511 | 0.090 | 13 |

Sail trawlers were of a round bilge shape and were classified as keel vessels (smacks) and flatbottom vessels (sloop, bomschuit). Smacks were streamlined vessels with a low Beam-Length overall [LOA]ratio (0.260). Bomschuiten were poorly streamlined vessels with a high Beam-LOA ratio (0.50). Sloops had an intermediate Beam-LOA ratio and a low Draft–LOA ratio (0.073). Keel vessels were dependent on harbours while flatbottom vessels could be hauled directly onto the beach. Sail surface area did not differ among vessel types and scaled to the square of vessel length (Figure 4c).

#### 2.3.2. Trawling practice

Beam trawling was the dominant fishing activity, although some sail vessels alternated beam trawling with, for instance, drift netting for herring or line fishing for gadoids. No trawling was possible when wind and tide were in opposite directions (Holdsworth, 1874; Holdsworth, 1883). The trawl was towed for a full tide or shorter when the border of a fishing ground was reached, the net got full or got stuck on the seafloor. Towards the turn of the tide, the trawl was hauled aboard and the vessel sailed upwind to an appropriate starting location for the next haul (Holdsworth, 1874). The effective trawling time was limited to 10 hours per day, corresponding to two tides. In areas with weak tidal currents, vessels could make tows of up to 9 or even 12 hours (De Zuttere, 1909; March, 1953).

Sailing trawlers towed one beam trawl at a speed of 2 knots over the ground, corresponding to 0.5 – 1.5 knots above the tidal speed (Holdsworth, 1874). Bomschuiten towed two beam trawls at a lower towing speed with the vessel axis perpendicular to tide (Petrejus, 1954). In the early 19^th^ century, sail trawlers made day-trips or trips of up to a few days. Vessels equipped with a well, that could keep their catch alive, were able to make longer trips. In the 1840s, trip duration increased to 6-8 days when vessels began to use ice to preserve their catch. The introduction of the fleeting system in England, where a carrier vessel collected the daily catch for transport to port, made it possible to trawl distant grounds and extend trip duration to 6 weeks - 2 months (March, 1953). The fleeting system was the dominant mode of operation in Yarmouth, Hull, and Grimsby.

Anchor seiners can operate in light winds but do not operate during night-time or when visibility is poor. Seiners typically make 8 hauls a day and require 1-2 hours to deploy and retrieve the gear (Blegvad, c[1950]). The SA per haul is dependent on the rope length. The rope length, that varies with fishing area and depth, increased from 300–500m around 1888 to 1000–2500m in 1929 (Mortensen and Strubberg, 1935).

Steam trawlers were no longer dependent on wind and tide. They trawled ∼20 hours per day when at the trawling ground (Masterman, 1914) with a towing speed of 2.5–3 knots (Currie et al., 2019). The increase in engine power of steam trawlers allowed vessels to use otter trawls with a horizontal net opening of 16 – 18.3 m.

#### 2.3.3. Days at sea

Only limited fishery statistics are available on the number of days at sea (das) and the effective trawling time per das (Table 3). Sail trawlers from Lowestoft and Ramsgate spend on average 209 das (sd=25) in the period 1906–1912 (Masterman, 1914). The effective trawling time of a sail trawler, which is dependent on the proportion of time required for sailing between the port and fishing ground, has an upper boundary of 10 hours.

**Table 3.** Overview of the mean (sd) of time budget parameters of sailing trawlers and steam trawlers based on effort statistics.

| Country | Days at sea<br>(year<sup>-1</sup>) | Trawling<br>hours<br>(das<sup>-1</sup>) | Comment | Source |
| --- | --- | --- | --- | --- |
| Sailing trawlers |  |  |  |  |
| England | 209 (25) |  | Lowestoft and Ramsgate<br>1906-12 | Masterman (1914) |
| Denmark | 65 – 100 |  | Sail seiners | Ministry of Agriculture (1897–<br>1915) |
|  | 150-200 |  | Motor seiners |  |
| Steam trawlers |  |  |  |  |
| England | 223 (16) | 15.8 | Fleet 1906-12 | Masterman (1914) |
| Netherlands | 235 (26) |  | Fleet | Anon (1913) |
| Germany | 199 (39) | 16.6 | Fleet | Herwig (1905)<br>Schnakenbeck (1927) |
| Germany | 283 | 16.8 (2.6) | One vessel 1904 | Herwig (1905) |

For Danish netters and seiners, the annual number of landing days in Esbjerg was reported for the period 1897-1914 (Ministry of Agriculture, 1897–1915) assumed to be representative of the Danish North Sea fleet. The number of days ranged between 65–100 days during the period when the fleet was still dominated by sailing vessels (1897-1900) and increased to ca 130 between 1901-1908 and ca 200 between 1909-1914 when motor vessels entered the fleet. Because anchor seining was restricted to the period between March and November, the annual number of fishing days was assumed to be 75% of these values.

The average number of days at sea of steam trawler fleets ranged between 189-235 das taking account of vessels that were inactive during part of the year (Table 3). The proportion active vessels of the Dutch steam trawler fleet varied between 0.71 (April-Aug) and 0.8-0.9 (Sept-Feb) (Anon, 1913). A single German steam trawler, that was active during the entire year, reported a total number of 283 das. The effective trawling time, taking account of the transit time between port and fishing ground, ranged between 15.8-16.6 hours per das.

With the growth of steam trawling, an increasing number of vessels trawled outside the North Sea. The percentage steam trawl effort in the North Sea dropped in England from 86% in 1903 to 55% in the 1910’s (Masterman, 1914), and in Germany from 96% in 1893 to 38-51% between 1899 – 1906 (Herwig, 1906).

#### 2.3.4. Fishing grounds

Information on trawling grounds was obtained from published maps and from reports (Table 4). The size and locations of the trawling grounds were estimated by digitizing the polygons directly from the published maps, or by estimating the spatial polygon where the depth matched the reported depth range and geographic boundaries using the *sf* library in R (version 1.0-17). Geographic location, depth range, and, where available, the size of the ground are given in 10.5281/zenodo.22078260.

**Table 4.** Fishing grounds. Overview of data used to estimate size and location of the trawling grounds.

|  |  |  |  |
| --- | --- | --- | --- |
| Country | Time period | Data type | Data source |
| England | 19<sup>th</sup> century | location, depth, size | Olsen (1878, 1885) |
|  |  | map | Alward (1911) |
| Germany | 19<sup>th</sup> century | Location, depth | Darmer (1894) |
|  | 1900 | Map | Schnakenbeck (1927) |
|  | 1920 | Map | Schnakenbeck (1927) |
| Belgium | 19<sup>th</sup> century | Distance from coast and/or fishing station | Orban de Xivry (1892)<br>de Zuttere (1909) |
| Netherlands | 19<sup>th</sup> century | Depth, distance, location | Collegie voor de Zeevisscherijen (1857–1911) |
|  | Early 20<sup>th</sup> century | Depth, location, map | Schouten, (1942)<br>de Veen (2024) |
| Denmark | 1880s | Map | Drechsel (1890) |
|  | 1890s | Map | Ministry of Agriculture (1893-1894) |
|  | 1920s | Map | Mortensen and Strubberg (1935) |

### 2.4. Mechanics of sail trawling

#### 2.4.1. Aerodynamic and hydrodynamic forces

Sail trawling is possible when there is sufficient wind in an appropriate direction to generate the thrust force to overcome the resistance of vessel and gear. The wind speed threshold was estimated by modelling the mechanics of sailing which is a complex interplay of hydrodynamic and aerodynamic forces (Slooff, 2015; Larsson et al., 2022). Equally complex are the forces acting on a gear (Paschen et al., 2000). Nevertheless, at the heart of the mechanics lay a number of equations that can be used to obtain a first order estimate of the relevant metrics for the different types of sailing trawlers operating in the bottom trawl fishery in the study period. Our simplified approach considers forces in the horizontal plane only. Details are given in 10.5281/zenodo.21904478.

The aerodynamic force can be decomposed in a thrust force (*T*) in the sailing direction (X-axis) and a side force (*S*_*A*_) perpendicular to the sailing direction (Y-axis)

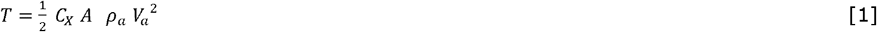

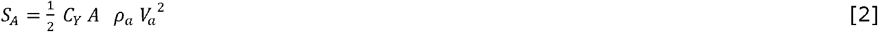

where *V*_*a*_ = apparent wind speed, *A* = sail surface area, *ρ*_*a*_ = air density, and *C*_*X*_ and *C*_*Y*_ are sail coefficients that vary with the angle *β* between the apparent wind and the vessel course. *C*_*X*_ gradually increase from negative values at low angles (*β* <15°) and reaches a peak *β*∼110°. *C*_*Y*_ increases to a peak at *β*∼35° and declines to zero at *β* =180°.

The aerodynamics side force *S*_*A*_ induces a hydrodynamic side force (*S*_*H*_) by the hull and keel with a component (*R*_*i*_) opposite to the thrust force

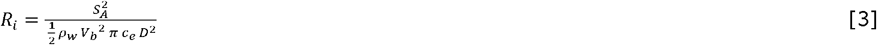

where *ρ*_*w*_ = water density, *V*_*b*_ = vessel speed in water,*c*_*e*_ = efficiency coefficient and *D* = draft of the hull.

When sailing at a constant speed, the aerodynamic forces are in equilibrium with the hydrodynamic forces. *T* will be equal to the sum of the resistance of the vessel (*R*_*vessel*_) and gear (*R*_*gear*_), and the aerodynamic side force will be equal to the hydrodynamic side force.

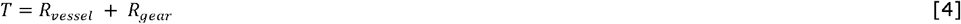

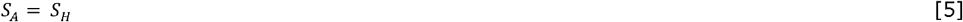

Vessel resistance (*R*_*vessel*_) comprised of the following components

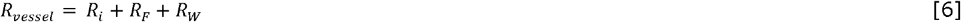

where *R*_*i*_ is the induced resistance; *R*_*F*_ is the friction of the water along the hull corrected for the effect of the hull shape on pressure differences; *R*_*w*_ is the wave making resistance, which is negligible at the towing speed and only becomes important when the speed approaches the maximum vessel speed.

Gear resistance *R*_*gear*_ is the sum of the hydrodynamic and geotechnical drag of its components (Paschen et al., 2000; O’Neill and Ivanović, 2016; Ghorai et al., 2025).

As only *T, S*_*a*_ and *R*_*i*_ are dependent on the apparent wind speed, *V*_*a*_ can be calculated by solving equation [4] for a given towing speed *V*_*b*_ and apparent wind angle *β* ∈ (0°-180°). The wind speed threshold (*WST*_*β*_) was obtained by converting *V*_*a*_ to the true wind speed and *β* to the true wind angle (*θ*) for a given *β*

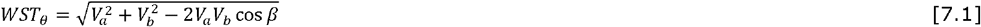

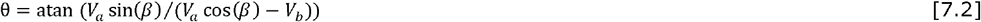

#### 2.4.2. Operational window for trawling

Trawling is possible when wind and tide are aligned (90° < *θ* < 180°). If we define *OR* as the operational range of wind directions with a valid *WST*_θ_, trawling will be restricted if θ_*min*_ > 90°.

The proportion of days favourable for trawling (*P*_*Ow*_) is given by

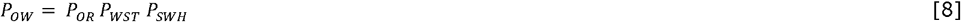

where *P*_*OR*_ = (180 − θ _*min*_)/90 for θ _*min*_ > 90°, *P*_*OR*_ = 1 for θ _*min*_ *≤* = 90°; *P*_*WST*_ is the proportion days when the wind speed >WST; *P*_*SWH*_ is the proportion of days when the significant wave height SWH<beam of the vessel.

### 2.5. Swept area estimation

#### 2.5.1. Swept area and SAR

The surface area swept annually (SA) was estimated as the sum of the *SA*_*f*_ of fleet *f*

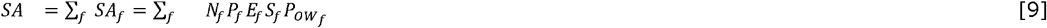

Where *N*_*f*_ = number of vessels, *P*_*f*_ = proportion of vessels engaged in bottom trawling, *E*_*f*_ = maximum number of days at sea (das) per year, and *S*_*f*_ = surface area swept by the gear per das (m^2^) and 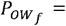 proportion of days with trawlable conditions.

For beam and otter trawlers *S*_*f*_ is calculated as *S*_*f*_ *= G*_*f*_*V*_*f*_*H*_*f*_, with *G*_*f*_ = width of the gear (m), *V*_*f*_ = the towing speed (m.s^-1^), *H*_*f*_ = number of trawling hours per das. For Danish seiners *S*_*f*_ was calculated as the product of the surface area swept during a single haul (Eigaard et al., 2016) and the number of hauls per das. The proportion of days with trawlable conditions 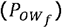 was estimated for each fleet following section 2.4.

Trawling intensity (SAR) is expressed as the ratio of the area swept (SA) over the surface area of the fishing grounds (A).

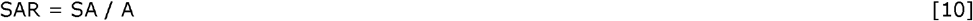

Confidence intervals of SA were estimated in a Monte Carlo simulation with 100 runs per fleet. Uncertainty was included for sail trawl fleets: *G*_*f*_ (CV=0.196), *E*_*f*_ (U(240,380)) and two levels of Ri (high, low); steam trawl fleets: *G*_*f*_ (CV=0.10), *E*_*f*_ (observed CV).

#### 2.5.2. Spatial distribution of SAR

The spatial distribution of SAR_tot_ of the international fleet was estimated by overlaying the SAR_f_ of fishing ground polygons of individual fleets assuming that SAR_f_ was evenly distributed in each fishing ground. SAR_f_ was assigned to the grid cells (1/30 degree latitude x 1/15 degree longitude, 2x2 nm at 60°N) falling within a polygon, and summed for each grid cell. The steam trawl effort of England and Scotland was distributed over the fishing grounds following the relative importance reported by Masterman (1914). The trawling footprint was calculated as the sum of the surface area of the fishing grounds at SAR>= 1 year^-1^ and the sum of the swept areas of fishing grounds trawled at SAR < 1 year^-1^ (Eigaard et al., 2017).

#### 2.5.3. Sensitivity analysis

The sensitivity analysis explored how a ±10% change in beam trawl width, towing speed, or vessel size affected SA of four fleets (Humber, Thames, Zijde, Ewer). The parameters were selected as they have a direct effect on the fished area and an indirect effect via the operational window. The selected fleets covered the range of vessel types and tidal speeds.

## 3. Results

### 3.1. Bottom trawling fleets

Around 1825, ca. 800 sailing vessels were bottom trawling at least part of the year. Their numbers increased, in particular after 1850, and reached a peak of 3500 sail trawlers in the 1880s (Figure 5a). Since then, their number decreased while the number of steam trawlers increased quickly. The expansion of bottom trawling was particularly pronounced in England that contributed ca two-thirds of the sail trawlers in the 1880s and an even larger proportion of the steam trawlers (Figure 5c). Information on individual fleets is provided in 10.5281/zenodo.22078260. The grounds of English vessels expanded over the entire North Sea down to 100m, whereas sail trawlers from Belgium, Netherlands, Germany and Denmark were mainly confined to grounds along the continental coast (Figure 5b,d).

**Figure 5.**
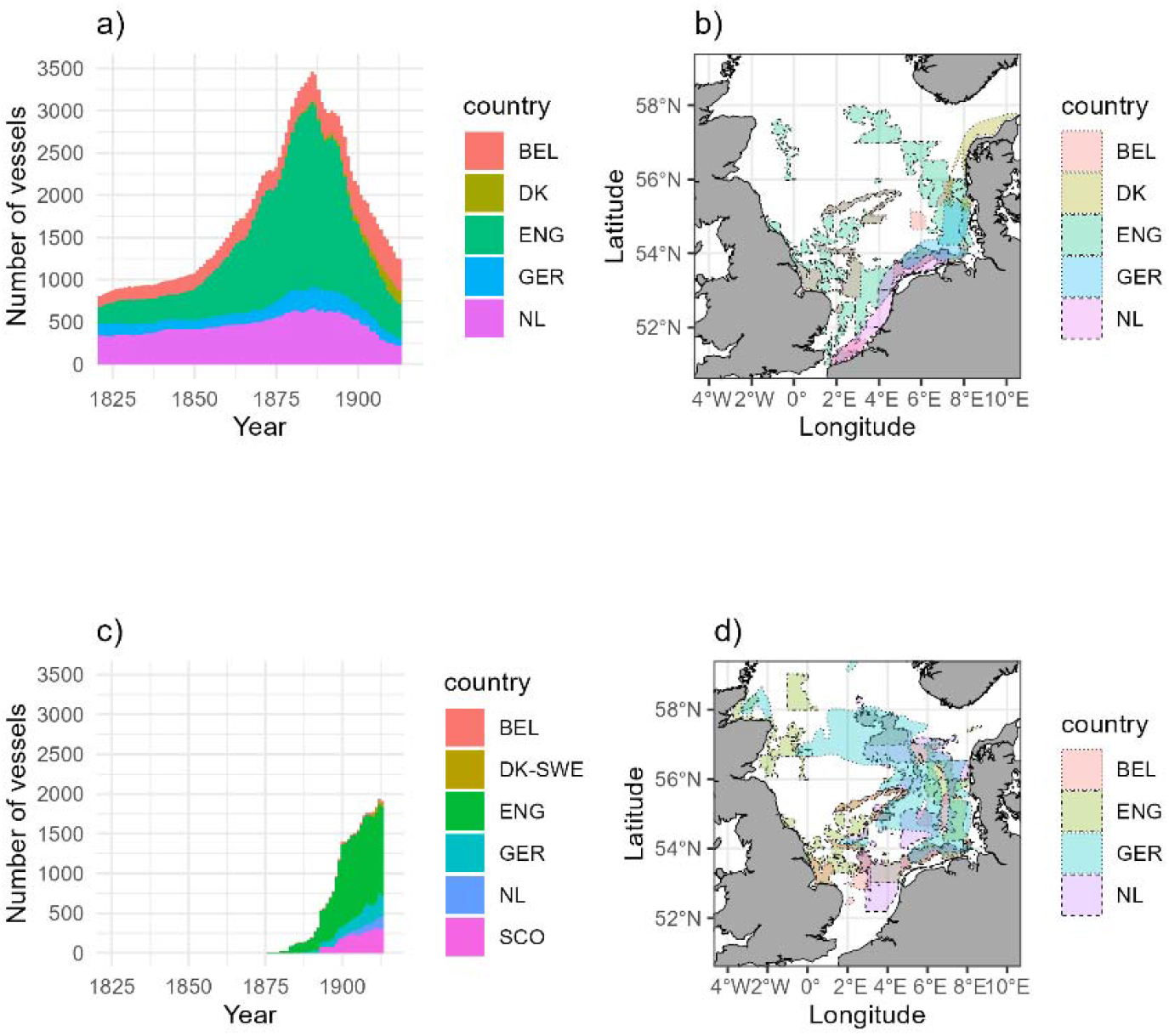
Temporal evolution of the number of (a) sailing vessels and (c) steam trawlers participating in the North Sea bottom trawl fishery and their respective fishing grounds (b, d).

### 3.2. Operational window

#### 3.2.1. Reference smack

The mechanics of sail trawling was quantified for a smack that is representative for the 1860s (Figure 6). The contour plot shows that the thrust force is largest when the angle between the true wind and vessel course is *θ* ∼90° (beam reach). The wind speed threshold (WST) at which the aerodynamic thrust force is in equilibrium with the resistance of the vessel and gear is determined by the resistance of gear and vessel and is lowest at broad reach (*θ*∼130°). Sailing closer to the wind will increase the induced resistance Ri and, as a consequence, will increase WST. Trawling opportunity parameters were estimated at *P*_*OR*_ = 0.70 and *WST*_*OR*_ =5.71 *m. s*^−l^ (Table 5). These estimates corresponded to the high gear and vessel resistance, which were based on information that a smack sailing at 8-9 knots would lose 6-7 knots when towing a beam trawl (Holdsworth, 1883). Trawling opportunity parameters for the low gear drag estimate, corresponding to the sum of the hydrodynamic and geotechnical drag of the gear components, were estimated at *P*_*OR*_ *=* 0.82 and *WST*_*OR*_ =4.72 *m. s*^−l^.

**Table 5.** Mean wind speed threshold (*WST*_*OR*_, m.s^-1^) and operational window parameters for a reference smack for different combinations of gear drag and induced vessel resistance (Ri). *WST*_*OR*_ was estimated over the operational range (*OR = 180° θ* _*min*_).

| Gear drag | $R_i$ | $WST_{OR}$ ($\text{m.s}^{-1}$) | Operational window | |
| --- | --- | --- | --- | --- |
| | | | $\theta_{min}$ | $P_{OR} = OR/90^\circ$ |
| Low (2.4 kN) | Low | 4.72 | 106 | 0.82 |
|  | High | 4.84 | 114 | 0.73 |
| High (3.4 kN) | Low | 5.59 | 109 | 0.78 |
|  | High | 5.71 | 117 | 0.70 |

**Figure 6.**
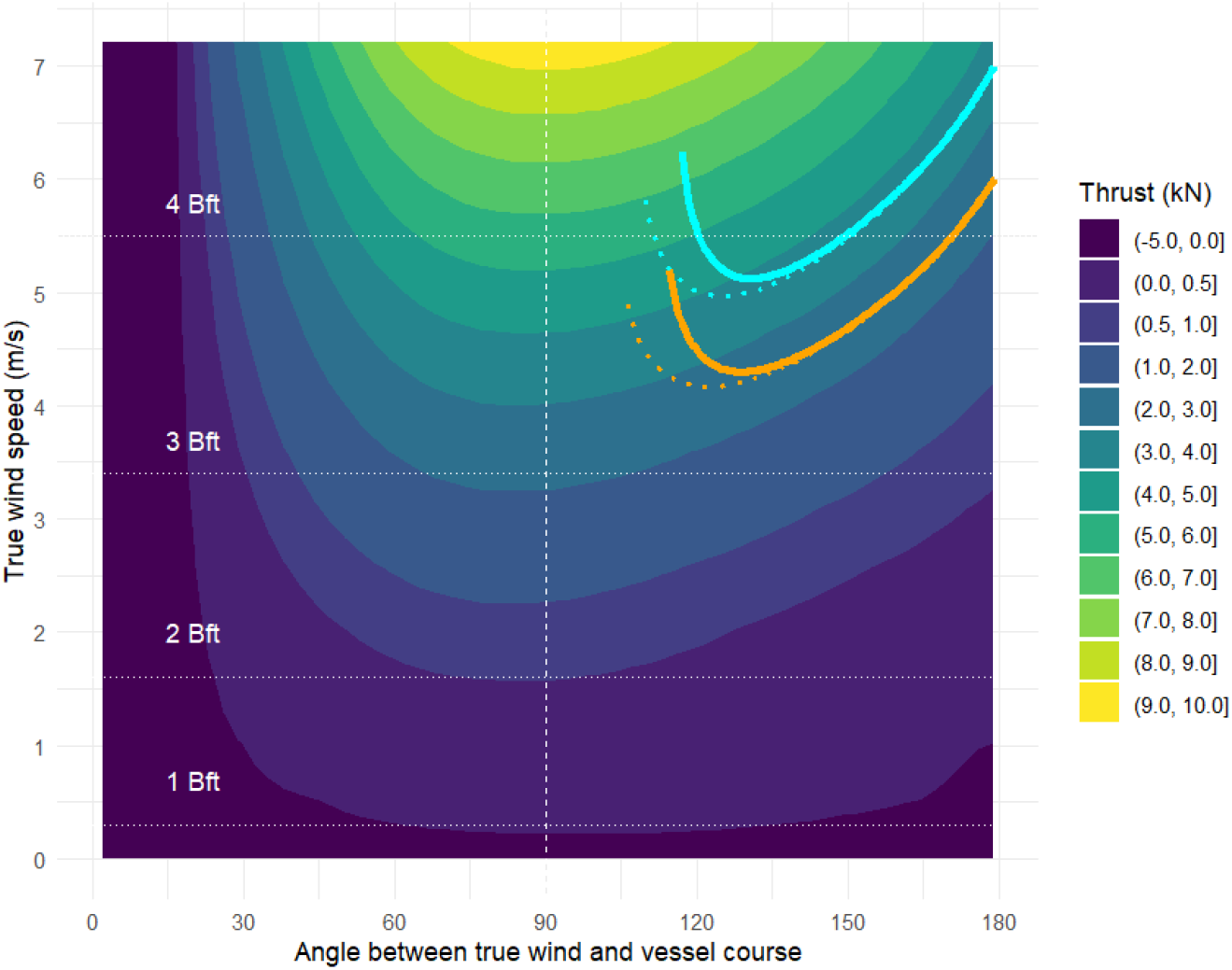
Contour plot of the thrust force (kN) in relation to the true wind speed (m/s) and the angle (*θ*, degrees) between the true wind and vessel course, and the wind threshold curves for a 20m smack towing a 12m beam trawl at 2 knot with the tide of 1 knot. Threshold curves are estimated for a high (cyan) and low gear drag (orange) and for a high (solid line) and low Ri scenario (dotted line). The vertical dashed line at 90° shows the boundary of the full operational range 90°-180°

#### 3.2.2. International fleets

*WST*_*OR*_ for individual fleets, taking account of differences in hull characteristics, vessel size, gear size, and tidal strength, ranged between 2-7 ms^-1^ (Figure7a). A high threshold was found for bomschuiten (NL_Zijde) that towed their gear with the vessel axis perpendicular to tide. Low thresholds were found for fleets operating in areas with high tidal speed (ENG_Thames, BEL_small and NL_Zeeland,). *WST*_*OR*_ was insensitive for uncertainty in Ri, in contrast to the operational range (OR) (Figure 7b). The OR was found to be restrictive for all fleets irrespective of Ri (*θ*_*min*_ > 90°).

**Figure 7.**
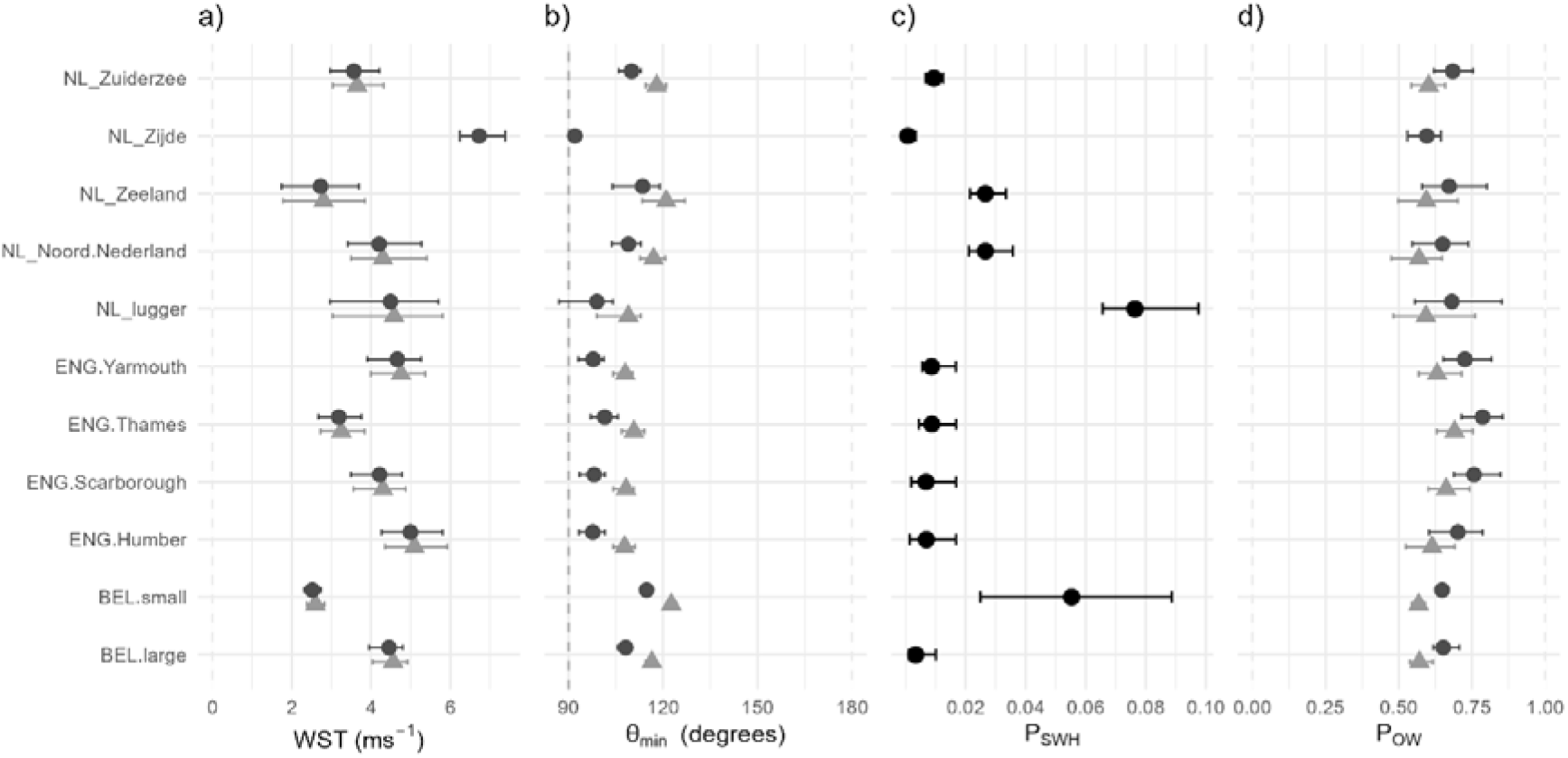
Median and 95% confidence interval of (a) wind speed threshold (WST); (b) minimum angle between wind and vessel course where trawling is possible (*θ*_min_); (c) proportion of days where the significant wave height exceeds the beam of the vessel (*P*_*SWH*_); (d) the proportion of favourable days for trawling (*P*_*Ow*_). Confidence intervals reflect the effect of 20% variation in the beam trawl length for a low (•) and high (Δ) Ri scenario. Confidence intervals in (c) represent the interannual variability in SWH

With the exception of small vessel fleets BEL_small and NL_lugger (*P*_*SWH*_ = 0.08-0.10), trawling opportunities were hardly affected by sea state (*P*_*SWH*_ <0.02) (Figure 7c).

The proportion of days with favourable trawling conditions (*P*_*Ow*_) varied between 0.64-0.78 for the low Ri and between 0.56-0.69 for the high Ri (Figure 7d).

### 3.3. Swept area

The SA of the international sail trawler fleet showed a strong increase from ca 35,000 km^2^ in the 1820s to a peak of ca 200,000 km^2^ in the 1880s and increased further to ca 500,000 km^2^ following the sharp increase in steam trawling (Figure 8a). The surface area of fishing grounds increased fourfold to ca 125,000 km^2^ in the 1880s and to 230,000 km^2^ in the 1910s. The mean trawling intensity SAR of the fishing grounds varied between 1-2 year^-1^. In the first half of the century, SAR decreased from 1.7 to 0.9 year^-1^ concurrent with the expansion of fishing grounds. In the second half, the increase in SA surpassed the expansion of fishing grounds and SAR increased to ca 1.75 year^-1^ at the heyday of sail trawling. After the arrival of steam trawlers SAR increased to ca 2 year^-1^ (Figure 8b).

**Figure 8.**
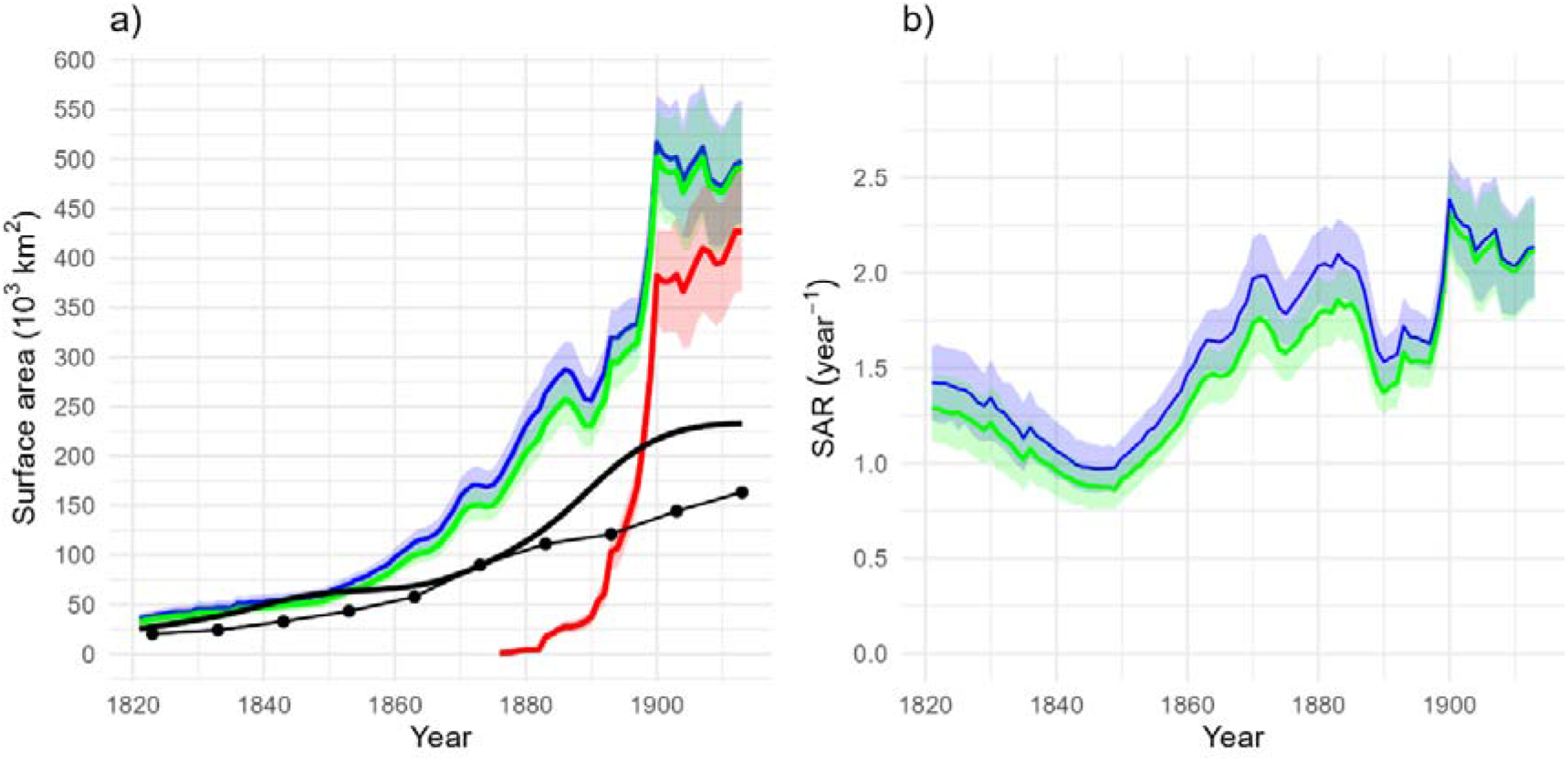
International bottom trawl effort: a) swept area (mean and 95% confidence interval, 10^3^ km^2^); b) swept area ratio (SAR, year^-1^). Green polygon = high Ri. Blue polygon = low Ri. Red polygon: steam trawlers. Black line = surface area of the fishing grounds (heavy black line). Dotted line (•) = trawling footprint.

Around 1830 intensive bottom trawling (SAR = 1-3 year^-1^) occurred in local grounds off the coast of England and Belgium. Grounds along the coast of Holland and Germany were trawled at SAR<1 year^-1^ (Figure 9a). In 1880, trawling had expanded over the central North Sea. Most grounds were trawled at SAR=1-2 year^-1^ with peak values of 4-5 year^-1^ (Figure 9b). In 1910, bottom trawling had expanded further, in particular north of 56°N, and several grounds in the central North Sea were trawled at SAR=5-7 year^-1^ (Figure 9c). A large part of the new grounds in the central and northern North Sea were trawled at SAR < 1 year^-1^.

**Figure 9.**
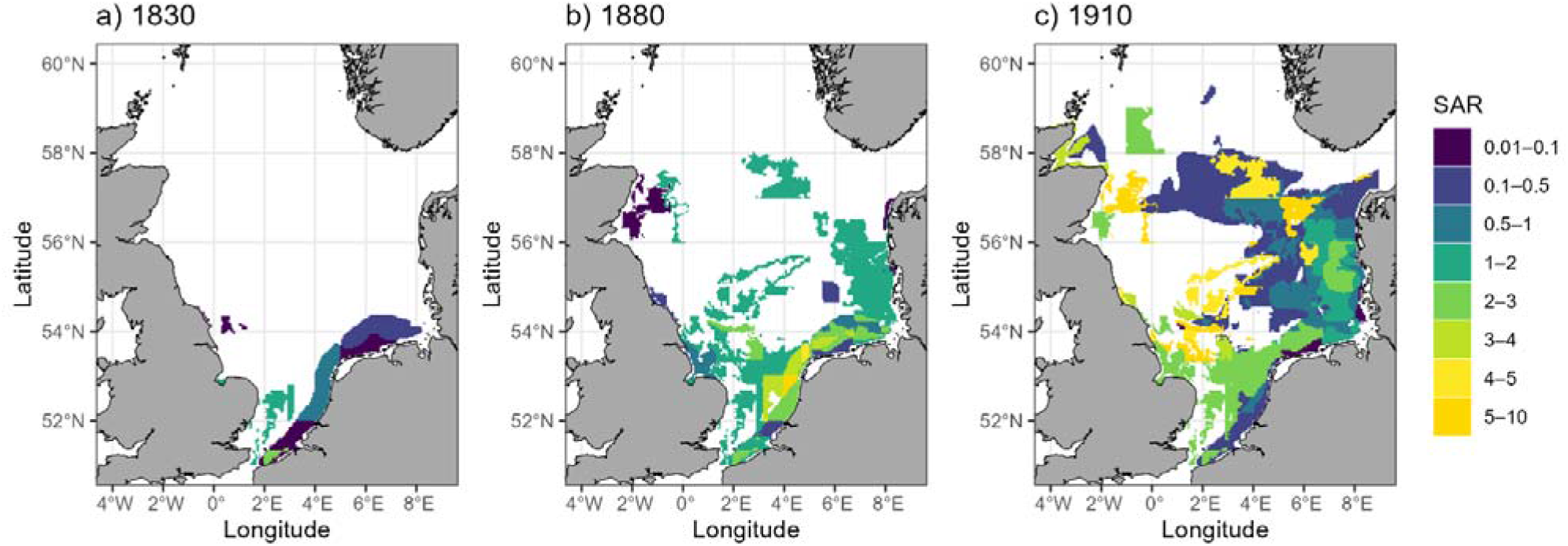
Swept area ratio (SAR, year^-1^) of the international bottom trawlers in 1830, 1880 and 1910.

The trawling footprint, measuring the unique seabed surface trawled per year, increased from ca 20,000 km^2^ in the 1820s to ca 160,000 km^-2^ in the 1910s (Figure 8a) covering on average 70% (sd=13%) of the surface area of fishing grounds.

Bottom trawling was concentrated in relative shallow water with a sandy sea floor (Table 6). Trawling depth increased gradually from 17–38 m in 1830 to 22-86 m in 1910. Concomitant with the increase in depth, trawling expanded into fishing grounds with lower sea bed stress. Sediment characteristics did not change much during the sail trawling era but with the advent of steam trawlers bottom trawling muddier sediments were trawled.

**Table 6.** Sea bed characteristics of trawling grounds: mean, 10^th^, 50^th^ and 90^th^ percentile of depth (m), sand (%), mud (%) and gravel (%) and seabed shear stress (N.m^-2^) weighted over the SAR by grid cell.

| Year | mean | p10 | p50 | p90 |
| --- | --- | --- | --- | --- |
| Depth (m) |  |  |  |  |
| 1830 | 28.2 | 17.1 | 30.0 | 37.6 |
| 1880 | 32.0 | 19.8 | 29.8 | 53.4 |
| 1910 | 49.0 | 22.0 | 39.0 | 86.0 |
| Sand (%) |  |  |  |  |
| 1830 | 90.0 | 73.6 | 94.6 | 99.5 |
| 1880 | 92.8 | 82.4 | 96.5 | 99.4 |
| 1910 | 88.3 | 68.8 | 94.6 | 98.9 |
| Mud (%) |  |  |  |  |
| 1830 | 3.2 | 0.3 | 1.7 | 7.5 |
| 1880 | 3.6 | 0.3 | 1.8 | 9.1 |
| 1910 | 7.3 | 0.4 | 2.1 | 19.1 |
| Gravel (%) |  |  |  |  |
| 1830 | 6.8 | 0.0 | 1.7 | 24.1 |
| 1880 | 3.6 | 0.0 | 0.2 | 13. |
| 1910 | 4.4 | 0.0 | 0.3 | 14.1 |
| Sea bed stress (N.m<sup>-2</sup>) |  |  |  |  |
| 1830 | 1.12 | 0.40 | 1.12 | 1.80 |
| 1880 | 0.49 | 0.06 | 0.37 | 1.13 |
| 1910 | 0.31 | 0.04 | 0.16 | 0.79 |

**Table 7.** Comparison of model estimates of trawling resistance, wind speed threshold (WST) and trawling opportunity window (POW) for low and high Ri with literature information. The range in model estimates reflect the variation (20%) in beam trawl size.

| Metric | Fleet | Gear drag | Model estimate |  | Reported value literature | unit | Source |
| --- | --- | --- | --- | --- | --- | --- | --- |
|  |  |  | Min Ri | Max Ri |  |  |  |
| Trawling resistance | Smack |  | 2.4 (1.2-4.1) |  | 3.4 (2.8-4.3) | kN | 1) |
| $WST_{OR}$ | Smack | low | 4.5 | 4.9 | 8.0 – 10.7 | $m.s^{-1}$ | 1) |
|  |  | high | 5.4 | 5.9 |  |  |  |
| $WST_{OR}$ | Sloop NL | | 3.1 – 5.2 | 3.3 – 5.8 | 5.5 – 7.9 | $m.s^{-1}$ | 2) |
| $P_{OR}$ | Belgian sloop | | 0.79 - 0.88 | 0.38 – 0.59 | 0.5 | | 3) |
| $P_{OR} * 280$ | Grimsby smack | | 202 - 233 | 150 - 200 | 137 | das | 4) |
| $P_{OR} * 280$ | Thames smack | | 241 - 260 | 164 - 215 | 209 (sd=25) | das | 5) |

1)Holdsworth (1874); 2) Venema (1869); 3) de Zuttere (1909); 4) March (1953); 5) Masterman (1914)

### 3.4. Sensitivity analysis

The sensitivity of the SA estimate varied between the input parameters and fishing fleets (Figure SM7.1 in 10.5281/zenodo.22078260). A 10% change in vessel size had an equally large effect on SA irrespective of fleet. The sensitivity for towing speed or beam trawl width differs among fleets. For the Thames fleet, SA is robust to changes in towing speed. In contrast, for bomschuiten (Zijde SKN) an increase in towing speed resulted in a decrease in SA. The Humber and German sailing fleets were intermediate. A change in beam trawl width results in a slightly smaller change (8%) in SA of the Thames and Zijde fleets, and a substantially smaller change for the Humber fleet (<=2%).

The difference in sensitivity between the fleets is related to the interaction between the direct effect of towing speed on SA and the higher wind force required to tow the gear at the higher speed. The latter will reduce the proportion of days with favourable winds (trawling opportunity window). The degree to which this will be reduced will depend on the vessel dimensions and strength of the tidal currents at the fishing grounds.

## 4. Discussion

This study reconstructs North Sea bottom trawl effort before the start of the collection of standardized fisheries statistics around 1900 (Watson and Tidd, 2018), extending the time horizon to the beginning of the 19^th^ century, and covering the heydays of sail trawling and early steam trawling. At that time, bottom trawling was rather insignificant compared to the driftnet fishery for herring and the hook and line fishery for gadoids (Schnakenbeck, 1927; Mortensen and Strubberg, 1931; Tesch and de Veen, 1933; Starkey et al., 2000; Lescrauwaet et al., 2013). After the end of the Napoleonic war (1815), bottom trawling showed a massive increase. The number of sail trawlers increased by a factor of 4.5 from ca 800 vessels in the 1820s to ca 3500 in the 1880s at the height of the sailing era. Coinciding with the increase in vessel size and gear size, the SA of sail trawlers increased 6 sixfold. With the rise of steam trawlers, SA increased twofold to almost 500,000 km^2^ and the area of fishing grounds increased to 245,000 km^2^.

As sailing vessels depend on wind and tide to trawl, SA is heavily dependent on the proportion of days with favourable wind conditions (*P*_*OR*_). To estimate *P*_*OR*_, we developed a model based on the fundamental equations of the hydro- and aerodynamic forces that can be parameterised with data on the hull and sail geometry of representative vessels. The model suggested that *P*_*OR*_ differed among fleets ranging between 55% – 80% of the days at sea. Given the simplifying model assumptions and the limited historical data available for parameterisation, results should be regarded as first-order approximations.

Uncertainty in the SA estimation was explored by taking account of the reported variability in beam trawl width (BTL) and the uncertainty in the number of days that fishing vessel can go out fishing (*Ef*). The effect of variability in BTL on SA will, to some degree, be compensated by the indirect effect on gear resistance and *P*_*OR*_ as illustrated in the sensitivity analysis. We assumed *Ef* to be uniformly distributed between 240 and 320 days. This concurs with (Holdsworth, 1883) reporting that sailing smacks spend at most nine months at sea, and information that the crew is at home for six weeks (March, 1953). Masterman (1914) reported that the Thames fleet spent 209 days at sea year^-1^ (sd = 25), but this value is based on effort statistics of the fleet which includes smacks that may have been inactive during part of the year. The *Ef* of Danish anchor seiners and steam trawlers was based on effort statistics. The reported number of landing days of the anchor seiners already excludes the days with unfavorable weather conditions and reflects that seining is only efficient in the warmer and lighter months of the year (April to November).

Despite the substantial margin of uncertainty surrounding the annual SA estimate, the upward trend is unmistakable. The trend, which is primarily determined by the growth in the number of vessels and their size, is robust and provides a sound basis for estimating the potential impact of trawling on the ecosystem.

The surface area of the trawling grounds were estimated by reconstructing fishing ground polygons. Fishing grounds of German and Danish fishers were based on published maps. Fishing grounds of Dutch and Belgian fishers were based on reported geographic or depth boundaries. English fishing grounds, and grounds of Dutch steam trawlers, were estimated from more detailed information on the geographic location, depth range and sometimes their size (Olsen, 1878; Olsen, 1885; de Veen, 2024). We infer that the latter grounds represent core fishing grounds were most, but not all, trawling activities took place. Recent studies indeed showed that bottom trawling is typically concentrated in the core part of the fishing grounds (Eigaard et al., 2017; Amoroso et al., 2018). The more general maps will likely reflect both core and peripheral areas.

The average trawling intensity of sail trawlers varied between SAR = 0.9-1.8 year^-1^, increasing to just over 2.0 year^-1^ with the rise of steam trawling. The trawling footprint, measuring the surface area trawled at SAR >= 1 year^-1^ increased to ca 150,000 km^2^ in the 1910s, covering on average 70% of the surface area of the fishing grounds. The reconstructed bottom trawl effort did not include the intensive trawl fishery for oysters that decimated the previous important North Sea oyster grounds (Bennema et al., 2020), and the shrimp fishery that occurred in English bays (Buckland and Walpole, 1878-9) and along the shallow continental coast of Belgium (De Zuttere, 1909).

The trawling intensity maps are based on the combined polygons of individual fishing grounds by fleet assuming a uniform effort distribution^1^. As fishing effort probably decrease with distance from port, the fishing effort in the more distant grounds may have been overestimated. The effect will be small for fleets that only trawled grounds close to port or the vessels participating in the fleeting system. A similar reconstruction by McLaverty et al. (2026), based on largely different historical sources and methodological approaches, estimated approximately 300,000 km^2^ of annual swept area for the British and Irish fleets during the early 20^th^ century, compared with approximately 500,000 km^2^ estimated here for the international North Sea fleet. The similarity in these estimates lends confidence to the reconstructed magnitude of historical trawling effort. In addition to this, the reconstructed SAR maps also broadly align, despite differences in how effort was spatially allocated.

The reconstructed bottom trawl effort provides a spatial explicit baseline for future studies of the impact of bottom trawling on the North Sea ecosystem, such as a study on the impact on the population dynamics of exploitation species. Plaice would be a good model species as it was one of the target species (Masterman, 1914). The annual harvest rate increased from 26% in 1892 to ca 50% in the period 1905 - 1914 corresponding to a twofold increase in SA of the international bottom trawlers (Rijnsdorp and Millner, 1996).

To assess the impact of the 19^th^ century bottom trawling activities on the sea floor and the benthos, a comparison is made with the impact assessment of the contemporary fishery. Table 8 summarises recently developed indicators for different time periods (Eigaard et al., 2017; Hiddink et al., 2017; Pitcher et al., 2017; Rijnsdorp et al., 2020). Nineteenth century bottom trawl effort increased 10fold and by 1900, fishing grounds covered almost 50% of the seafloor down to 200m depth. A century later, trawl effort had further increased by 30% and fishing grounds covered almost the entire sea floor area. The surface area of sea floor that was trawled annually increased from 5% in 1830 to almost 30% in 1900, and just over 60% in 2010. Historic trawling intensities varied between SAR = 1 and 2 year^-1^ and were at the end of the 19^th^ century higher than in 2010.

**Table 8.** Trawling impact indicators of the international bottom trawling fleets in the North Sea (0-200m) in the period 1820 – 2010. Fishing ground surface and trawling footprint are expressed as a percentage of the surface area between 0-200m depth.

| Indicator | 1830 | 1880 | 1900 | 2010 |
| --- | --- | --- | --- | --- |
| Fishing ground surface (%) | 7.2% | 25.0% | 46.0% | 93% <sup>(1)</sup> |
| Trawling footprint (%) | 4.8% | 22.3% | 28.9% | 63% <sup>(1)</sup> |
| Swept area (1000 km<sup>2</sup>) | 41 | 201 | 499 | 648 <sup>(1)</sup> |
| SAR (year<sup>-1</sup>) | 1.1 | 1.6 | 2.2 | 1.25 <sup>(1)</sup> |
| RBS | 0.970 | 0.948 | 0.926 | 0.87 <sup>(2)</sup> |

Sources: 1) Eigaard et al. (2017); 2) Rijnsdorp et al. (2020)

Bottom trawling damage and kill organisms in the path of the trawl. Recently, a simple method to quantify the trawling impact has been developed (Pitcher et al., 2017) which estimates the relative benthic status (RBS) given the rate of exposure to trawling (SAR), the depletion rate due to a trawling event and the recovery rate of the organisms in the community. The method has been applied to several present-day fisheries including the bottom trawl fleets in the North Sea (Rijnsdorp et al., 2020; Pitcher et al., 2022). To estimate the impact of historic bottom trawling we used the same methodology and parameterization but only adjusted the depletion rate imposed by the historic beam trawl. Because historic beam trawls were relatively light without tickler chains, we assumed a depletion rate=0.026, similar to a contemporary otter trawl targeting cod and plaice, but much lower than the depletion rate of contemporary beam trawls (d=0.14). Bottom trawling in 1830 reduced the RBS within the trawling footprint by 3%. Trawling impact more than doubled by the turn of the century (7%). Contemporary bottom trawling reduced the RBS by In 2010 the impact almost doubled to 13%. Because the proportion of long lived organisms in the benthic community likely decreased over time, the historic impact may have been underestimated.

Although the historic beam trawls were relatively light, the shoes and ground rope will have impacted epifaunal structures or biogenic material, which are long lived species with a high sensitivity. Rich epifaunal communities occur predominantly in areas with a hard bottom. Although hard bottom habitats pose a threat of damaging or losing the gear, fishers likely entered these areas either accidentally, or deliberately in pursuit of a good catch, and gradually changed the nature of the habitat. Fisheries information on the location of foul areas, compiled in the 1880s (Olsen, 1878; Olsen, 1883; Darmer, 1894), showed that 8.1% of the surface area of the North Sea (5-100 m) was comprised of rocks and stones, 0.5% comprised of large pieces of peat (moorlog) and 5.9% comprised of dense oyster beds (10.5281/zenodo.22078260). In a map of 1920, the extent of the sea floor considered untrawlable (foul to very foul) was reduced to 3.6% and none of the areas with moorlog or oysters were shown (Close, 1920). Although the introduction of bobbins in early 20^th^ century may have enabled bottom trawlers to move into foul grounds, the reduction in the extent of untrawlable ground could be related to a change in the seafloor due both the intensive bottom trawling and the fishery for oysters (Bennema et al., 2020),

Bottom trawling may also impact the carbon storage of the seafloor through mixing and resuspension of sediment (Epstein et al., 2022; Hiddink et al., 2023). Sediment resuspension rate is a function of the silt fraction of the sediment and the hydrodynamic drag of the gear (O’Neill and Ivanović, 2016; Rijnsdorp et al., 2021; van der Reijden et al., 2025). The transition from sail to steam trawling will have accelerated sediment resuspension due to the increase in towing speed and expansion of trawling grounds. The increase in towing speed will have resulted in a threefold increase in hydrodynamic drag and a twofold increase in the mean silt content of the trawling grounds. With the tenfold increase in SA, the total amount of sediment resuspended will have increased by about 60, in broad agreement with the 30-fold increase estimated for the English and Irish bottom trawl fleets (McLaverty et al., 2026).

## Supporting information

SupportingMaterialPart1_Part2

## Author Contributions

ADR.: Conceptualization, methodology, software, validation, formal analysis, investigation, data – curation, writing – draft, writing – editing and reviewing, visualization. FPB: Investigation, data – curation, writing – editing and reviewing. FV: writing –reviewing. ORE: writing – editing and reviewing. JAT: Investigation, data – curation, writing – editing and reviewing. CM: Conceptualization, methodology, investigation, writing – editing and reviewing

## Acknowledgements

We gratefully acknowledge the help and critical feedback on trawling history from A. Klok (Wageningen University), dr F. Loomeier and on the aero- and hydrodynamics of sail trawling from drs Wick Hillege (Dykstra-NA), J. Hernandez Montfort, M. Verhulst (MARIN), G. Jacobi (Delft-University), prof dr Ana Ivanovich, dr F. O’Neill and prof dr J. van Leeuwen (Wageningen University).

## Data Availability Statement

Data to reproduce the results of this paper are available in 10.5281/zenodo.22078260.

## Supporting information

Supporting information is available in 10.5281/zenodo.22078260 providing information of beam trawl gears, characteristics of sailing trawlers, reconstruction of the trawling grounds, estimation of the number of sail trawlers and steam trawlers, input parameters for the estimation of the swept area of individual fleets, and the results of the sensitivity analysis. A detailed presentation of the mechanics of sail trawling is presented in 10.5281/zenodo.21904478.

## Footnotes

1 Only for English steam trawlers information was available on the spatial distribution of fishing effort over individual fishing grounds (Masterman, 1914).

