## SupportingMaterialPart1_Part2 for "From sailing to steam trawling: the evolution of bottom trawl effort in the North Sea"

Part 1. **10.5281/zenodo.22078260**

[1)]Wageningen Marine Research, PO Box 68, 1970 AB IJmuiden, The Netherlands; [2)]MarHis, Haren, The Netherlands; [3)]VFC Veenstra fishery consultancies, Haarlem, The Netherlands; [4)]National Institute of Aquatic Resources (DTU Aqua), Technical University of Denmark, DK-2800 Kongens Lyngby, Denmark; [5)]Centre for Ecology and Conservation, University of Exeter, Penryn Campus, Cornwall, TR10 9FE, England.

#### 1. Abstract

This repository presents supporting material for the paper "From sailing to steam trawling: the evolution of bottom trawl effort in the North Sea", in which a reconstruction of the fishing effort of the international bottom trawl fleets in the North Sea between 1820 and 1914 is presented. Historical data on gear dimensions, sailing vessel specifications, and vessel numbers, are presented for different fleets. Input data to reconstruct fishing grounds by country and fleet are presented, as well as the spatial polygons. Operational characteristics are presented required for the estimation of the surface of the seafloor swept by each fleet. Finally, results of a sensitivity analysis of the swept area estimation is presented.

### 2. Fishing gear.

North Sea sail trawlers predominantly used a beam trawl to catch bottom dwelling fish. A compilation of data on the dimensions of beam trawls is given in ***SM1.tables.xls/beam.trawl.length***

A detailed description of the beam trawl gear of the Grimsby smack "Willy and Ada" was reported by (Collins, 1889). The vessel towed a single 14 m beam trawl (Table 2.1). Dimensions of the net are presented in Table 2.2 and Figure 2.1. The surface area of the twine was estimated at 31.7 m$^2$ interpreting that the reported (square) mesh size correspond to the distance between two adjacent knots. The stretched mesh size then corresponds to 2M. No data were reported on the twine diameter $d$ and the number of meshes $N_d$. We assumed a single braided twine diameter of 4 mm slightly larger than the twine diameter of beam trawl nets used in the contemporary fishery. $N_d$ was estimated as $N_d$=$P_d/M_v$, where $P_d$ = depth of the panel, $M_v$ is the vertical width of the mesh. $M_v$ can be calculated as $M_v$ = $2M cos(\alpha/2)$, where $\alpha = asin(M_h/2M)$ and $M_h$ is the horizontal width of a mesh $M_h$= $P_w/N_t$, and $P_w$ = width of the panel. The diameter of double braided twine was taken as 1.64 times the twine diameter.

Collins (1889) further noted that the length of ground rope used by the sailing vessels varied across fishing stations. The stations in the north generally used a ground rope where the distance between the beam and the centre of the rope was roughly equal to the width of the beam trawl. In southern stations, shorter ground ropes were deployed with a distance between the beam and the centre of the ground rope being 4/5 of the beam trawl.

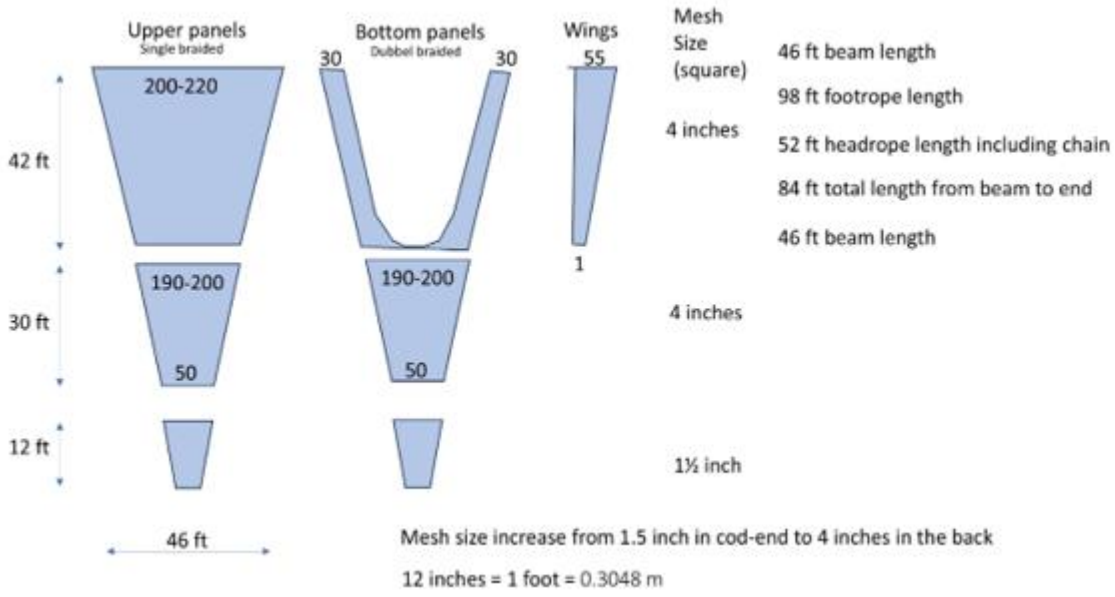

*Figure 2.1. Net drawing and dimensions of the beam trawl net of the Grimsby smack "Willie and Ada" as provided by Collins (1889). Values in the net panels show the number of meshes on the top and bottom row of the panels. The number of meshes along the length of the panels was estimated from the panel length and the mesh size.*

Information on the beam trawl gear used by Dutch fishers is available from the publications of (Venema, 1869; Petrejus, 1954; de Veen, 2024). Bomschuiten trawled two beam trawls of about 8 m width (Collegie voor de Zeevisscherijen, 1854). The length of the net was 13 m with a codend braided of 'very strong twine' (hemp).

De Veen (2024) described dimensions of a beam trawl used by the schokkers and botters participating in the beam trawl fishery in the coastal waters. The beam trawl length was 8 m. Nets were made of hemp ('hennep omslag' of 250-350) and the cod-end was double braided. The length of the net and cod-end was 12 and 2 m, respectively and the mesh size was reduced from 0.07 in the front part to 0.05 in the aft part. Table 2.3 shows the number of meshes of the net panels assuming that the reported mesh size reflects the bar size. The surface area of the netting was estimated at 7.0 $m^2$.

Venema (1869) describes the fishery in the province of Groningen in the northern part of the Netherlands and provided quantitative information on the vessel and beam trawl gear used (Table 2.3). A sailing schuit from Zoutkamp of 12.6 m (LOA) deployed a 8.6 m beam trawl with a net and cod-end length of 15.3 and 5.1 m, respectively, and a ground rope of 10.3 m. The central part of the ground rope was over a distance of 1.8 m deployed with lead. Assuming that the vessel used a similar mesh size than reported by de Veen (2024), the surface area of the twine was estimated at 13.4 $m^2$. We interpreted the mesh size to represent the stretch mesh size (March, 1953).

The available information on the beam trawl gear and netting used by sail trawlers is summarised in Table 2.5 and complemented with the data reported for a Scottish steam trawler by Fishery Board for Scotland (1894).

Table 2.1. Dimensions of gear components of the Grimsby trawling smack "Willie and Ada" (Collins, 1889).

| Gear component | | Value | unit | Value | unit |
|---|---|---|---|---|---|
| Beam trawl | Length | 46 | feet | 14.0 | m |
| | Diameter | 8 | Inch | 0.20 | m |
| | Frontal surface | | | 2.8 | $m^{-2}$ |
| | Weight | | | 1050 | $kg.m^{-3}$ |
| Trawl head (2x) | Weight | 180 | Pound | 81.6 | kg |
| | Height | 4 | Feet | 1.22 | m |
| | Breath | 2 10/12 | Feet | 0.86 | m |
| | Width | 4 | inch | 0.10 | m |
| | Frontal surface | | | 0.124 | $m^{-2}$ |
| | Footprint surface | | | 0.088 | $m^{-2}$ |
| Foot rope | Length | 98 | feet | 29.9 | m |
| | Diameter | 7 | Inch | 0.17 | m |
| | Weight | | | 21 | $kg.m^{-1}$ |
| Head rope | Diameter | 2.5 | Inch | 0.06 | m |
| Netting | Twine surface area | | | 31.7 | $m^2$ |

Table 2.2. Dimensions of the netting panels of a 14 m beam trawl net of the Grimsby smack "Willie and Ada" made of tarred hemp (Collins, 1889).

| Net panel | Number of meshes | | | Mesh size[2)] (m) | Twine diameter (m) | Surface m2 |
|---|---|---|---|---|---|---|
| | Nt | Nb | Nd[1)] | | | |
| Top 1 | 210 | 195 | 63 | 0.2032 | 0.004 | 10.4 |
| Top 2 | 190 | 50 | 45 | 0.2032 | 0.004 | 4.4 |
| Wing | 55 | 1 | 63 | 0.2032 | 0.004 | 1.4 |
| Wing | 55 | 1 | 63 | 0.2032 | 0.004 | 1.4 |
| Bottom 1 | 60 | 60 | 63 | 0.2032 | 0.0065[3)] | 5.0 |
| Bottom 2 | 190 | 50 | 45 | 0.2032 | 0.0065[3)] | 7.1 |
| Cod-end (round) | 100 | 64 | 48 | 0.0760 | 0.0065[3)] | 1.9 |

1) Nd was estimated from the reported length of the net panels. 2) Stretched mesh size. 3) The diameter of double braided twine was taken as 1.64 times the twine diameter.

Table 2.3. Reconstructed characteristics of the netting panels of a 8 m beam trawl net (de Veen, 2024)

| Net panel | Length (m) | Width (m) | Number of meshes | | | Mesh size[2)] (m) | Twine diameter (m) | Surface netting (m2) |
|---|---|---|---|---|---|---|---|---|
| | | | Nt | Nb | Nd | | | |
| 1 | 2.5 | 8.0 | 171 | 139 | 18 | 0.14 | 0.003 | 1*1.2 |
| 2 | 7.5 | 6.5 | 163 | 50 | 63 | 0.12 | 0.003 | 2*2.4 |
| 3 | 2.0 | 2.0 | 60 | 24 | 20 | 0.1 | 0.003 | 2*0.25 |
| Cod-end | 2.0 | 0.8 | 24 | 24 | 20 | 0.1 | 0.006 | 0.3 |
| Total | 14.0 | | | | | | | 7.0 |

Table 2.4. Reconstructed characteristics of the netting panels of a 8.6 m beam trawl net (Venema, 1869)

| Net panel | Length (m) | Width (m) | Number of meshes | | | Mesh size[2)] (m) | Twine diameter (m) | Surface netting (m2) |
|---|---|---|---|---|---|---|---|---|
| | | | Nt | Nb | Nd | | | |
| 1 | 1.5 | 8.6 | 184 | 172 | 11 | 0.14 | 0.003 | 1*0.8 |
| 2 | 9 | 8 | 201 | 116 | 75 | 0.12 | 0.003 | 2*4.3 |
| 3 | 5 | 4.7 | 140 | 26 | 50 | 0.1 | 0.003 | 2*1.2 |
| Cod-end | 5.1 | 0.9 | 26 | 26 | 51 | 0.1 | 0.006 | 2*0.8 |
| Total | 20.6 | | | | | | | 13.4 |

Table 2.5. Summary of the dimensions (m) of beam trawls

| Vessel type | Smack | Schokker | Schuit | Bomschuit | Steamer |
|---|---|---|---|---|---|
| Vessel length | 24 | 12.2 | 13.7 | 12.2 | |
| Beam trawl | 14 | 8 | 8.6 | 8 | 16.5 |
| Diameter beam | 0.20 | | 0.15 | | 0.55 |
| Trawl head | 1.22 | 0.8 | 0.9 | | 1.14-1.22 |
| Foot rope | 29.9 | 10-12 | 10.3 | | 37.8 |
| Head rope diameter | 0.06 | 0.06 | | | 0.20 |
| Net length | 25.6 | 12 | 15.3 | | 28.7 |
| Mesh size | 0.20–0. 076 | 0.07–0.05 | | | 0.076-0.051 |
| Cod-end length | 3.65 | 2 | 5.1 | | 7.3 |
| Cod-end mesh | 0.076 | 0.05 | | | 0.038 |
| Netting ($m^2$) | 31.7 | 7.0 | 13.4 | | |
| Trawling speed (knot) | 2 | 2 | 2 | 1.5 | 2.5 |
| Source | 1) | 2) | 3) | 4) | 5) |

1) Collins (1891); 2) de Veen (2024); 3) Venema (1869); 4) Collegie voor de Zeevisscherijen (1854); Petrejus (1954); 5) Fishery Board of Scotland (1894)

### 3. Vessel characteristics

The characteristics of the sailing vessels are compiled in ***SM1.tables.xls/Hull.data*** providing information on the surface area of the sails, length overall (LOA), length on the waterline (LWL), beam (maximum width) of the vessel, draught and tonnes of the vessel, and hull shape (keel, flatbottom) and vessel type (smack, sloop, bom).

Data on size (tonnage), year when built and station of registration was available for over 1000 English trawling smacks from (Olsen, 1878; Olsen, 1885; March, 1953). Average vessel size increased from around 30 tons in the beginning of the 19th century to about 70 tons in the mid-1870s and declining to about 40 tons in the early 20th century (Table 3.1). Vessel size in the northern stations (Humber: 75 ton in 1878; Scarborough:72 ton in 1878) was substantially larger than the size of the newly built smacks in the Yarmouth and Lowestoft (52 ton in 1878). The corresponding vessel length (LWL) increased from about 10 m at the start of the 19th century to close to around 15 m and 19 m at the end of the century in the southern stations (Thames) and Humber stations, respectively.

Table 3.1. England. Average size of newly built trawling smacks by fishing station and decade (data from Olsen, 1878 and March, 1953). The LOA was estimated from the glm model fitted through the paired recordings: log(LOA) = 1.70685 + 0.34259*log(Tonnage) (explained deviance = 0.810)

| Decade | Tonnage (100 feet$^3$) | | LOA (m) | | n |
|---|---|---|---|---|---|
| | mean | sd | mean | sd | |
| All stations England | | | | | |
| 1810 | 31.0 | | 13.9 | | 1 |
| 1820 | 15.0 | | 10.8 | | 1 |
| 1830 | 37.6 | 5.3 | 14.6 | 0.70 | 5 |
| 1840 | 42.3 | 7.3 | 15.2 | 0.86 | 9 |
| Thames (Ramsgate, Barking, Lowestoft) | | | | | |
| 1850 | 44.0 | 7.9 | 20.1 | 1.3 | 3 |
| 1860 | 45.3 | 6.1 | 20.4 | 1.0 | 32 |
| 1870 | 50.8 | 8.5 | 21.1 | 1.2 | 46 |
| 1880 | 51.1 | 15.2 | 21.1 | 2.5 | 12 |
| 1890 | 41.1 | 16.0 | 19.3 | 2.9 | 19 |
| 1900 | 43.1 | 13.2 | 19.8 | 2.2 | 29 |
| 1910 | 39.0 | 7.0 | 19.3 | 1.2 | 3 |
| 1920 | 40.0 | 7.0 | 19.4 | 1.3 | 9 |
| Yarmouth | | | | | |
| 1850 | 37.7 | 4.4 | 19.1 | 0.7 | 36 |
| 1860 | 41.2 | 4.7 | 19.7 | 0.8 | 139 |
| 1870 | 45.8 | 7.7 | 20.4 | 1.1 | 120 |
| 1880 | 62.3 | 11.5 | 22.6 | 1.5 | 3 |
| Scarborough | | | | | |
| 1850 | 47.1 | 7.9 | 20.6 | 1.2 | 9 |
| 1860 | 52.1 | 6.5 | 21.3 | 0.9 | 17 |
| 1870 | 64.5 | 10.7 | 22.9 | 1.3 | 20 |
| Humber (Hull, Grimsby) | | | | | |
| 1850 | 44.9 | 10.4 | 20.1 | 1.8 | 42 |
| 1860 | 56.0 | 8.4 | 21.8 | 1.1 | 232 |
| 1870 | 70.8 | 7.1 | 23.7 | 0.8 | 550 |
| 1880 | 63.3 | 31.4 | 22.3 | 4.4 | 3 |
| 1890 | 86.0 | | 25.4 | | 1 |

### 4. Trawling fleets

#### Sail trawlers

##### England.

Detailed accounts of the trawl fishery of sailing vessels is given by (March, 1953; Robinson, 2000). Beam trawling started off in the early 19th century when smacks from Brixham migrated seasonally (later permanently) to fishing ports in the Thames area. Vessels seasonally targeted flatfish with a beam trawl off the southern English east coast or off the Dutch coast, and cod and haddock off the Lincolnshire coast using the trawl or line. In the 1840s, the use of ice and the introduction of the fleeting system, where the daily catch a fleet of trawling smacks was transferred to a carrier vessel for transport to the fish market, enabled fishers to extend their fishing trips and exploit distant fishing grounds. As a result, the trawl fishers moved to other fishing ports that were more favorably located to the new fishing grounds.

In Lowestoft, no trawling took place until the mid-19th century when smacks from Ramsgate started to use the port. Smacks worked as single boaters making trips of 6-10 days in the southern North Sea (Masterman, 1914; March, 1953).

In Yarmouth, home to a large fleet of herring drifters, the first trawling smacks arrived in the 1840s. Their number increased during the 1850s, when trawling fleets from elsewhere began using the port because of its proximity to the fishing grounds. It was not until the early 1870s that local smacks were converted to allow them to target herring with drift nets during the autumn season (three months) and flatfish with a beam trawl during the remainder of the year (March, 1953).

The discovery of the Silver Pits caused a rush northwards, with smacks from Brixham and Ramsgate beginning to settle in Hull and Grimsby. Located close to the coalfields, supplying fuel for the steam carriers that had replaced sail carriers by the 1880s, and well connected to major consumer markets by railway, Grimsby and Hull became the centres of the trawl fishery.

In Scarborough, local fishers were drifting for herring and longlining for gadoids. The first trawling smacks settled in the early 1830s in summer and early autumn to trawl local grounds. The number of locally owned trawling smacks increased since the 1840s.

For most of the 19th century only anecdotal information is available on the number of 1st class sailing trawlers operating from the various fishing stations: ***SM1.tables.xls/n.sail.eng.various***. Annual numbers are reported from 1889 onwards by Garstang (1900): ***SM1.tables.xls/ n.sail.eng.garstang1910***; and Masterman (1914): ***SM1.tables.xls/n.sail.eng.masterman1914***.

For the purpose of our study, the English sailing trawlers were grouped into four bottom trawl fleets: Thames, Single boaters (Yarmouth), Fleeters (Yarmouth, Grimsby, Hull) and Scarborough. Fleet numbers were estimated for each year by linear interpolation of the number by fishing station (Figure 4.1) and summed for each fleet.

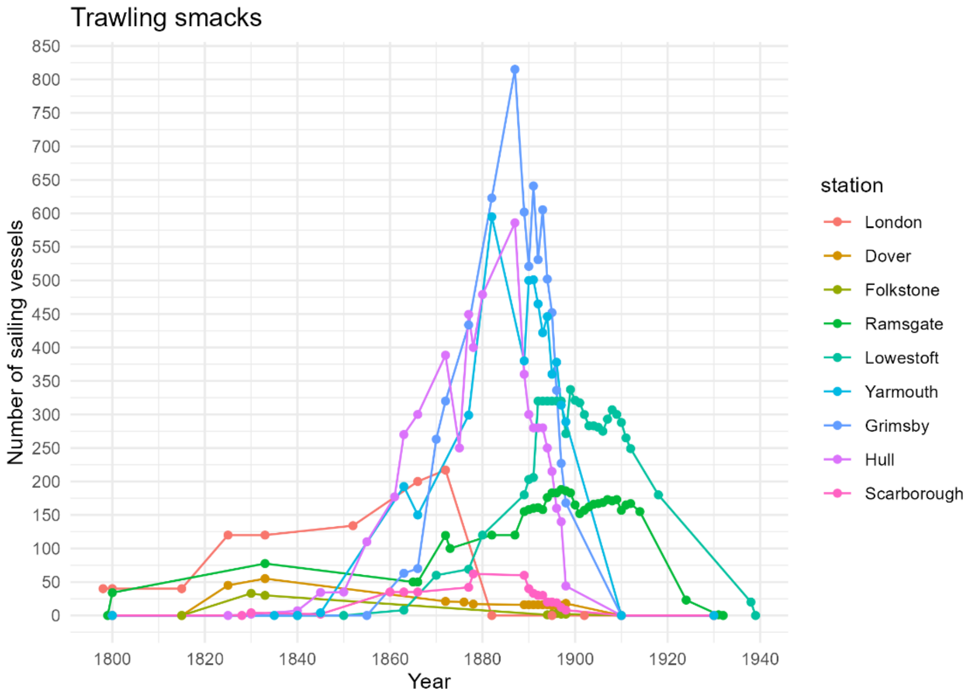

*Figure 4.1. England: number of smacks participating in the bottom trawl fishery in the North Sea.*

#### Belgium

A detailed account of the Belgium fisheries in the 19th century and early 20th century is given by (De Zuttere, 1909) based on an enquiry conducted in 1905. De Zuttere distinguished four different fisheries: 1) drift net fisheries for herring in the central North Sea (Grande Peche du Hareng); 2) hook and line fishery for cod in winter (North Sea) and summer (Iceland); 3) fishery for fresh fish using a variety of fishing gear; 4) brown shrimp fishery. For the reconstruction of the effort of the bottom trawl fishery, we focus on the fishery for fresh fish (Peche de Maree).

***SM1.tables.xls/n.sail.be*** presents the number of sailing vessels by fishing station that were engaged in the fishery for fresh fish (De Zuttere, 1909). The fishery for fresh fish comprised a variety of combinations of target species and fishing gear. Besides the beam-trawl fishery for flatfish, vessels targeted roundfish with hook and line, herring and sprat with drift nets, and demersal fish with stocknets or seine nets. Following its introduction in Ostend in 1822, the beam trawl gradually became the dominant fishing gear (Figure 4.2). Other fishing stations adopted beam trawling at a later stage. According to the Chambre des Représentants (1866), beam trawling was the main fishery throughout the year in Blankenberge at that time, and was also practised in Heyst, Ostend, and Nieuwpoort. In De Panne, fishers began beam trawling during the summer months in 1880. The larger vessels in Heyst, De Panne, Oostduinkerke, and Koksijde used the beam trawl for most of the year, except during the winter, when they switched to the coastal herring fishery. Vessels from Nieuwpoort trawled during the summer, while in winter they fished for cod on the Dogger Bank using hook and line. The cod fishery from this station ended in 1884.

For the purpose of our study, the Belgian sailing vessels were grouped into two bottom trawl fleets. A fleet of small vessels (<20 ton) of flat-bottomed vessels operating from the beach, and a fleet of large vessels (>20 ton) of keel vessels operating from Oostende throughout the time period and from Blankenberge since 1880. Table 4.1 shows the frequency distribution of vessel size for the different fishing station around

1900, which were used to allocate the number of vessels over the small and large categories used in the present study.

Table 4.1. Belgium. Size composition of the fishing vessels engaged in the fishery for fresh fish by fishing station around 1900 (de Zuttere, 1909).

| Station | Number of vessels by size class (tonnage, RT=2.83 m$^3$) | | | | |
|---|---|---|---|---|---|
| | Sailing vessels | | | | Steam trawler |
| | <5* | >20 | 10-20 | >20 | >20 |
| Oostende | 1 | 22 | 4 | 93 | 22 |
| Nieuwpoort | 7 | | 21 | 5 | |
| Blankenberge | | | 26 | 12 | |
| Heijst | | | 31 | 1 | |
| De Panne | 5 | | 18 | | |
| Oostduinkerke | 2 | | 20 | | |

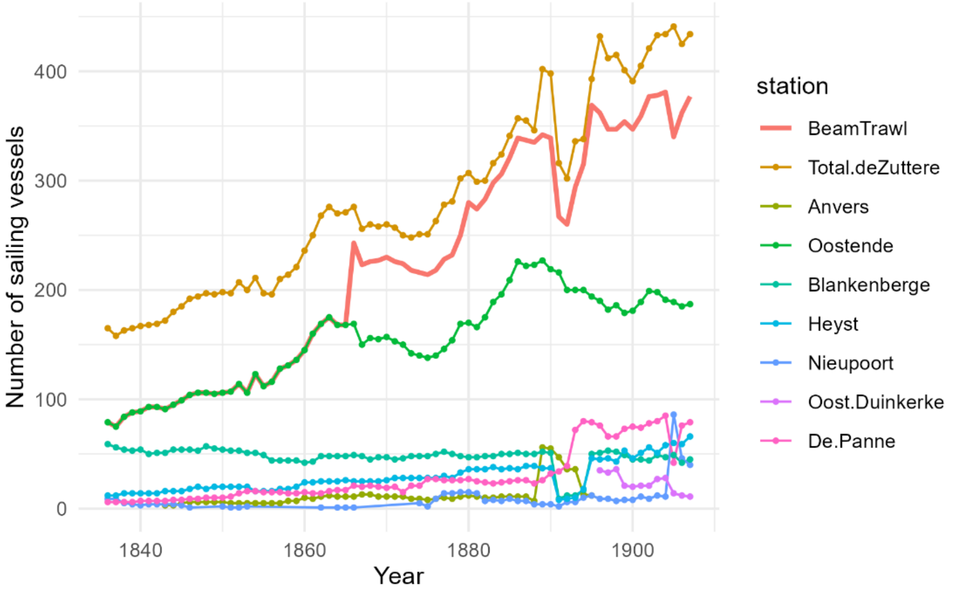

*Figure 4.2. Belgium: number of sailing vessels engaged in the fishery for fresh fish by fishing station according to de Zuttere (1909). Beam trawl shows the number of sailing vessels using a beam trawl.*

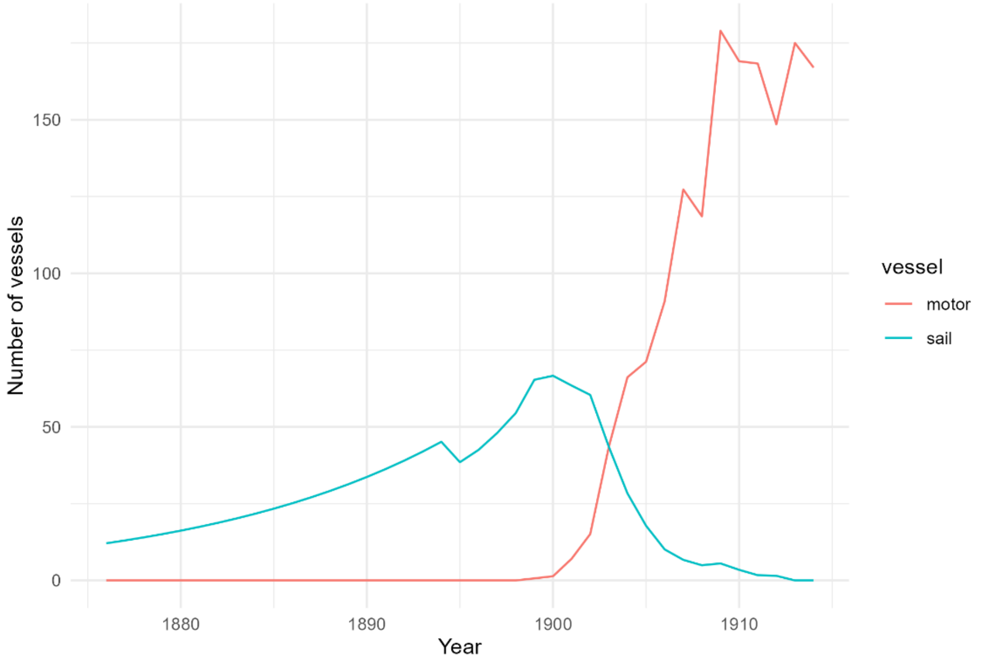

*Figure 4.3. Denmark: Number of fishing vessel engaged in the anchor seine fishery in the North Sea.*

#### Denmark

***SM1.tables.xls/n.anchorseine.dk*** presents the number of sailing and motor vessels anchor seining in the North Sea. Danish sailing vessels started bottom trawling using an anchor seine at the end of the 19$^{th}$ century (Figure 4.3). Just after the turn of the century, sailing vessels were replaced by motor vessels.

The 1895 to 1911 data originate from registered vessels along the western coast of Jutland in the official fisheries statistics of Denmark (Ministry of Agriculture, 1897-1915). All vessels categorised as cutters and 50% categorised as boats included. The 1912 to 1914 data originate from vessels reported practising their fishery in North Sea and Skagerrak in the official fisheries statistics of Denmark (Ministry of Agriculture, 1897–1915). All vessels categorised as vessels above 15 BRT and 50% categorised as 5-15 BRT included. The 1876 to 1894 data are predicted number of vessels from fitting the observed number of fishing vessels (1895 to 1911) to a nonlinear least-squares exponential model

#### Germany

A detailed account on the North Sea fishery (Hochseefischerei) of German sailing vessels is given by (Schnakenbeck, 1927). The fishery was conducted by fishers from Finkewardern and Blankenese using a beam trawl[1].

***SM1.tables.xls/n.sail.ger*** and Figure 4.4. presents the data on the number of sailing vessels (ewers) operating from Finkewardern and Blankenese in the period 1750-1927 (Schnakenbeck, 1927). For the study period, vessel numbers in missing years were estimated by linear estimation for each stations and summed.

[1] Prior to 1820, fishers have been using a static gear.

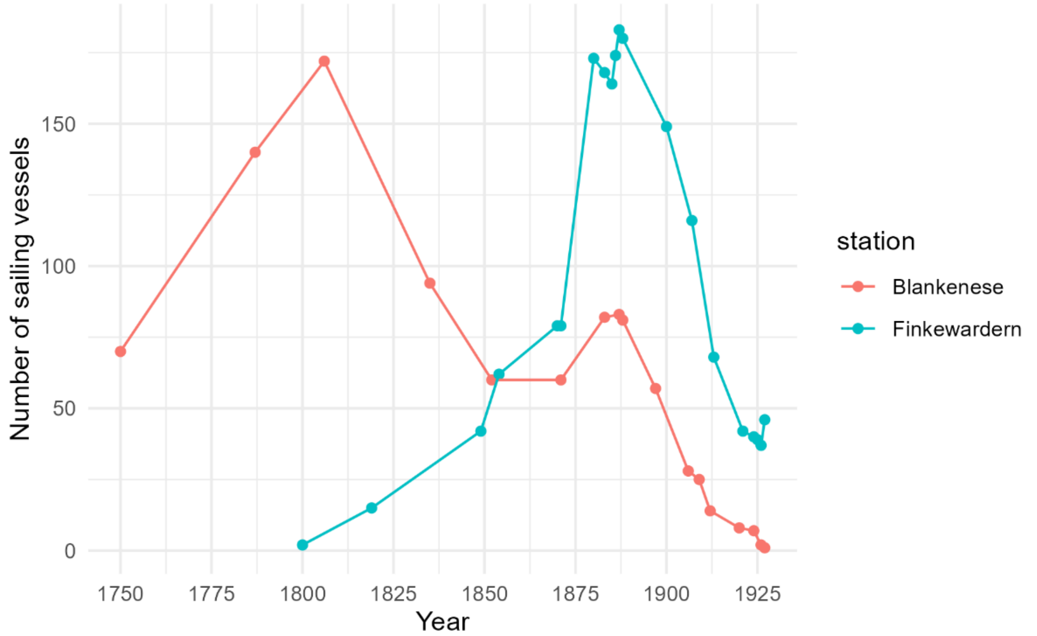

*Figure 4.4. Germany. Number of sailing vessels engaged in bottom trawling in the North Sea.*

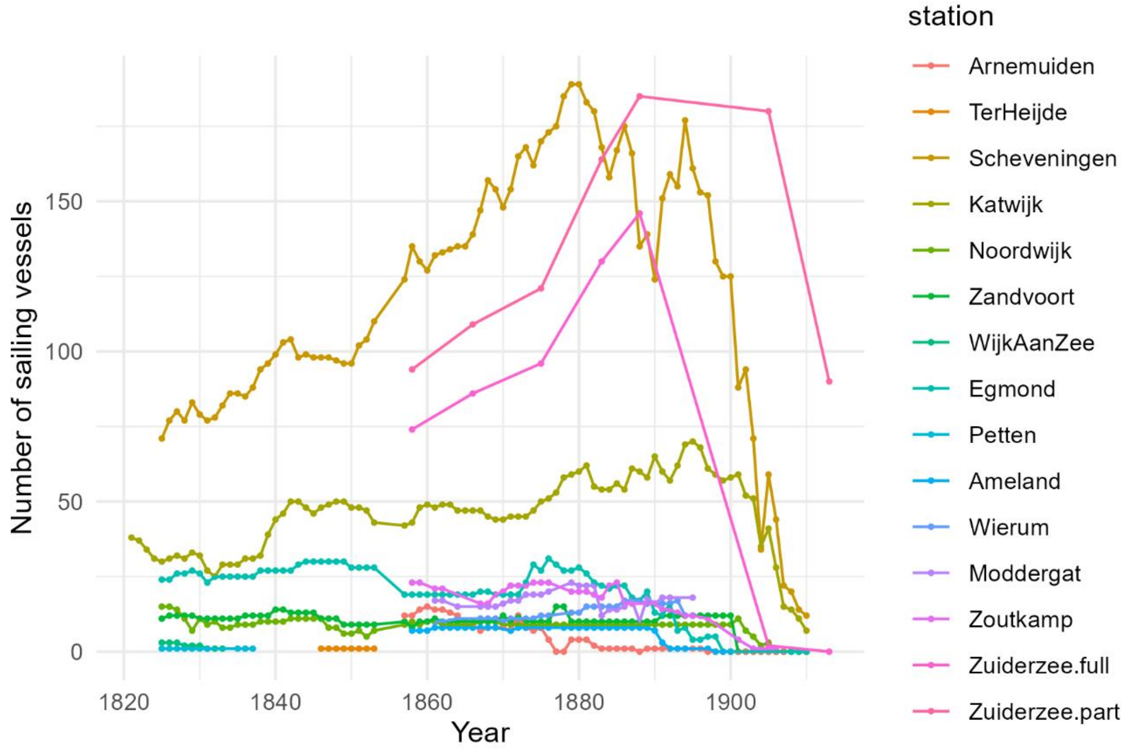

*Figure 4.5. Netherlands. Number of sailing vessels engaged in bottom trawling in the North Sea from various fishing stations.*

#### Netherlands.

The major fisheries in the Netherlands were the herring driftnet fishery and the cod hook-and-line fishery, while bottom trawling was of lesser importance (Beaujon, 1884). Bottom trawling was conducted from the villages along the coast of Holland (de Zijde), Zeeland, Friesland, Groningen and villages along the border of the Zuiderzee. The changes in the number of sailing vessels during the study period is shown in Figure 4.5.

Beam trawling was a seasonal activity of bomschuiten operating from the sandy beach along the coast of Holland. Beam trawling in spring was alternated with driftnet fishing for herring in autumn and hook and line fishing for gadoids in winter. Bomschuiten from Egmond were trawling throughout the year (le Compte, 1831; Beaujon, 1884). Trawling was not allowed between 15 November and 15 February (1 February since 1837) (Beaujon, 1884). Prior to 1857, the trawling season lasted till mid-August. After fisheries restrictions were lifted in 1857, bomschuiten started their herring season by mid-June and trawling was exercised the first five months of the year (Collegie voor de Zeevisscherijen, 1857-1911). The number of bomschuiten are presented by fishing station in ***SM1.tables.xls/n.sail.nl.kramer*** (Kramer, 1984) and ***SM1.tables.xls/n.sail.nl.zijde*** (Collegie voor de Zeevisscherijen, 1857–1911).

Another major fishery with beam trawls was carried out by fishers from Zuiderzee villages (Redeke, 1907; Dorleijn, 1982). Most of the Zuiderzee fishers restricted their activities to estuarine waters, but fishers from Urk (UK), Voldendam (VD), Enkhuizen (EH) and Huizen (HZ) fished in the coastal waters of the North Sea for flatfish with the beam trawl or roundfish using hook and line. Some vessels alternated the fishery in the North Sea with a fishery in the Zuiderzee for herring (January – April) and anchovy (May – July). The number of Zuiderzee vessels that were engaged in the North Sea fisheries was estimated by applying the proportion of North Sea vessels in 1889 to the fleet size estimated for the villages from where fishers were fishing in the North Sea (UK, VD, EH, HZ) in the years prior to 1889: ***SM1.tables.xls/n.sail.nl.zuiderzee*** (Redeke, 1907; De Zuttere, 1909; Dorleijn, 1982). The fleet size in 1820 was arbitrarily set at 75% of the fleet size in 1858. The proportion of time spent in North Sea activities was estimated at 8 month for the part time vessels and 12 month for the full time vessels. The median vessel size of the North Sea vessels was 24 tons (Redeke, 1907).

Limited bottom trawling took place from villages in the mouth of the Scheldt and Meuse (Zeeland) and along the Wadden coast. Fishers from Zeeland alternated between trawling for flatfish and hook and line fishing for gadoids. The number of vessels involved, and their size, was small: ***SM1.tables.xls/n.sail.nl.other*** (Collegie voor de Zeevisscherijen, 1857–1911). In the northern provinces, vessels seasonally fished for flatfish or gadoids (haddock). In the first half of the 19th century, vessels from Zoutkamp were dredging for oysters, but this fishery disappeared after the 1850s (Venema, 1869).

For the purpose of our study, the Netherlands sailing trawlers were grouped into regional groups: Zeeland, Holland, Zuiderzee (full), Zuiderzee (part), northern provinces (Friesland, Groningen). The annual number of sailing trawlers by fleet were estimated by linear interpolation of the number by fishing station (Figure 4.5) and summing the annual numbers over the stations by fleet.

#### Steam trawlers

The annual number of steam trawlers is presented in ***SM1.tables.xls/n.steamer***. Data on the English and Scottish fleet is based on (McLaverty et al., 2026). Data of other countries are from Bulletin Statistique (ICES, 1956).

### 5. Trawling grounds

Trawling grounds were based on reports describing the name, geographic location, depth range, and, where available, the size of the ground. Individual fishing grounds of English and Dutch bottom trawlers were reconstructed by estimating the polygon within geographic boundaries where depth matched the reported depth range.

#### England

***SM1.tables.xls/fg.olsen*** provides the depth range (fathom), geographic boundaries (degrees.minutes) and the year when the bottom trawling commenced of the individual grounds of English bottom trawlers reported by (Olsen, 1878; Olsen, 1883; Olsen, 1885). When the starting year was not reported, it was inferred from (Alward, 1911). Figure 5.1. shows a map of the reconstructed fishing grounds of the bottom trawl fleets and the start of bottom trawling (***fg.olsen.gpkg***).

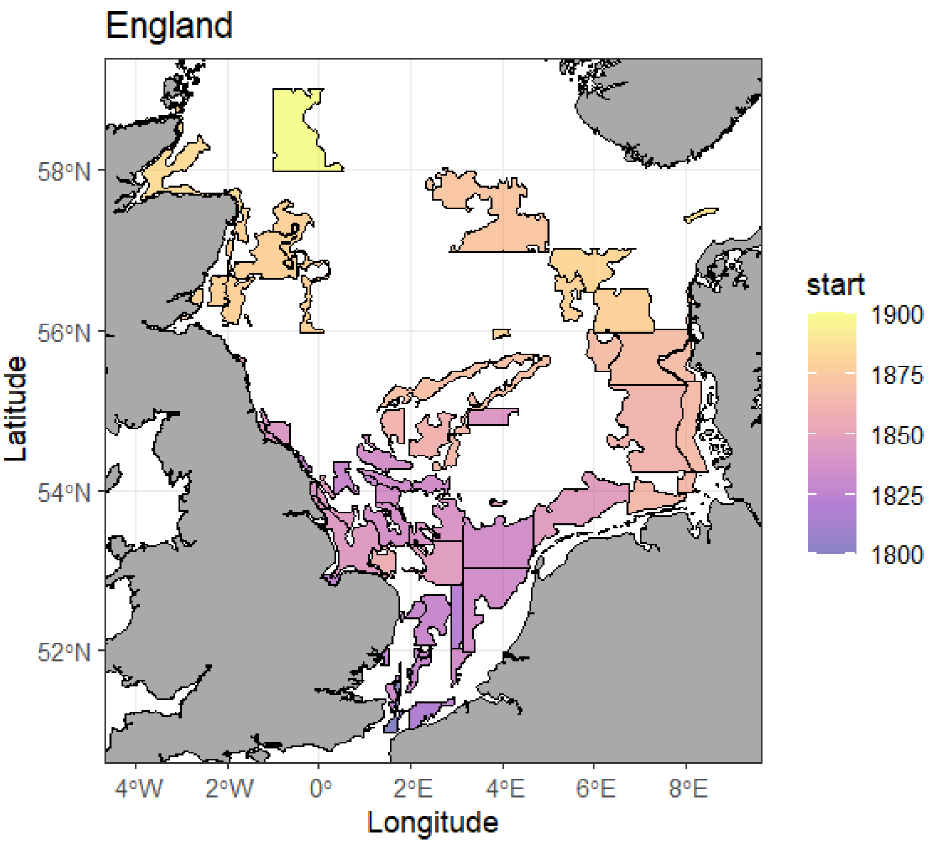

*Figure 5.1. Expansion of bottom trawling by English vessels during the 19th century based on the estimated fishing ground polygons.*

Fleet specific fishing grounds (***fg.fleet.eng.sco.gpkg***, Figure 5.2) were estimated by selecting a subset of individual grounds of Figure 5.1. The grounds were selected according the fishing patterns of the fishing stations described by March (1953). The smacks from the Thames stations (London, Ramsgate, Lowestoft) predominantly trawled in the southern North Sea (Figure 5.2a). Smacks from Yarmouth operated as single boaters (Figure 5.2b) or participated with the vessels from the Humber region (Grimsby, Hull) in the fleeting system (Figure 5.2c). Smacks from Scarborough mainly trawled in the western grounds (Figure 5.2d).  Steam trawlers mainly operated in the central and northern North Sea (Masterman, 1914)(Figure 5.2.ef).

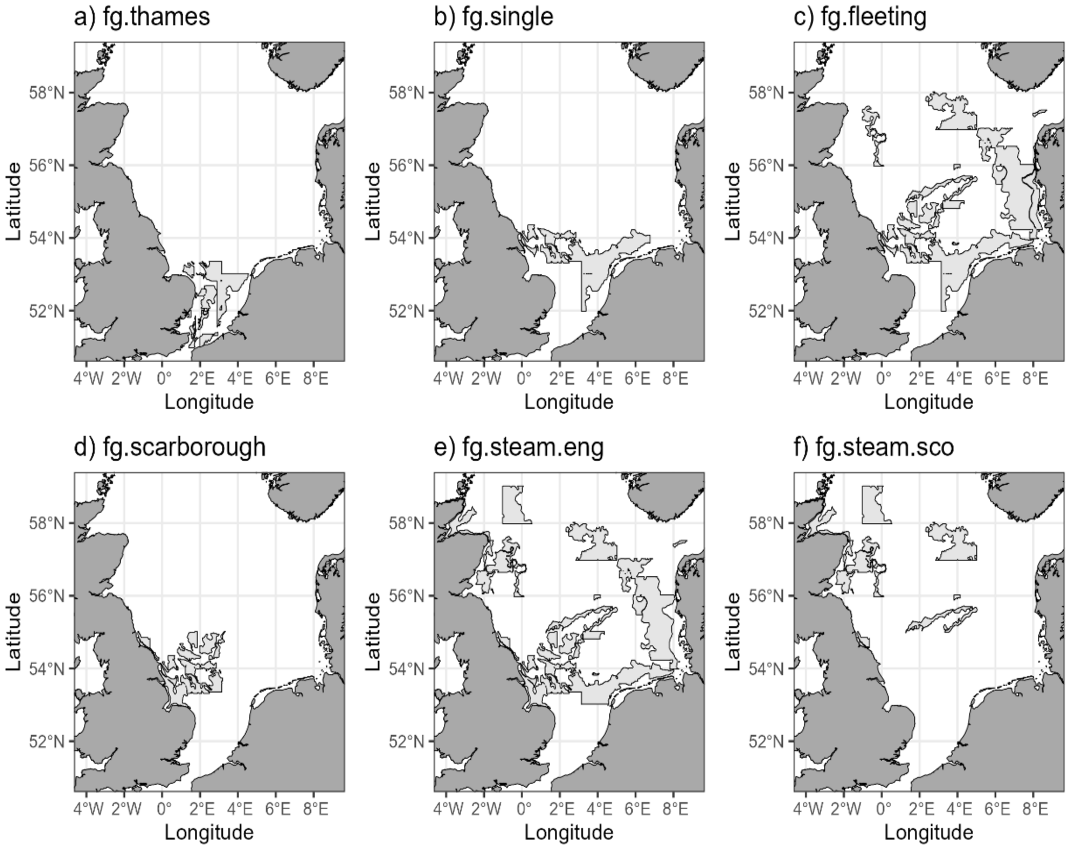

*Figure 5.2. England and Scotland. Fishing grounds of the different fleets of sail and steam bottom trawlers in 1900.*

#### Belgium

Fishing grounds of Belgian bottom trawlers are shown in Figure 5.3 (***fg.fleet.bel.gpkg***). According to de Zuttere (1909), small vessels (<20 ton) operated along the Belgian coast within 25 nm from their station (Figure 5.3a). Larger vessels (>= 20 ton) trawled off the Belgian-Dutch coast and off the English coast (Figure 5.3b), extending their grounds to the Dogger and White Bank at the end of the 19th century (Figure 5.3c). The larger vessels followed a distinct seasonal pattern (Orban de Xivry, 1892; Verbrugghe, 1932) (Table 5.1). Based on the frequency of mentions, the grounds of the highest importance are the grounds off the west Frisian islands (Terschelling (7), Ameland (5) and Borkum (5)); off England  (Silver Pits (6), Botney Ground (5)) and the White Bank (5) in the German Bight.

Table 5.1. Major fishing grounds of Belgian vessels (>20 ton) by month (Orban de Xivry, 1892)

| Month | Fishing ground |
|---|---|
| January | Ameland, Borkum, Terschelling |
| February | Borkum, Doggerbank, Ameland. |
| March | Borkum, Norderney, Heligoland, Dogger bank, Silver Pits, Botney ground, Wittebank |
| April | Doggerbank, Botney ground, Wittebank |
| May | Ameland, Terschelling |
| June | Hornreef, Ameland, Terschelling, Silverpits, Wittebank, Borkum |
| July | Silverpits, Terschelling, Smiths Knoll, Outer Dowsing, Leman, Owen |
| Agust | Silverpits, Wittebank, Botney ground |
| September | Botney ground, Claydeep, Silverpits |
| October | Wittebank, Silverpits, Claydeep, Terschelling |
| November | Terschelling, Ameland, Smiths Knoll |
| December | Borkum, Terschelling |

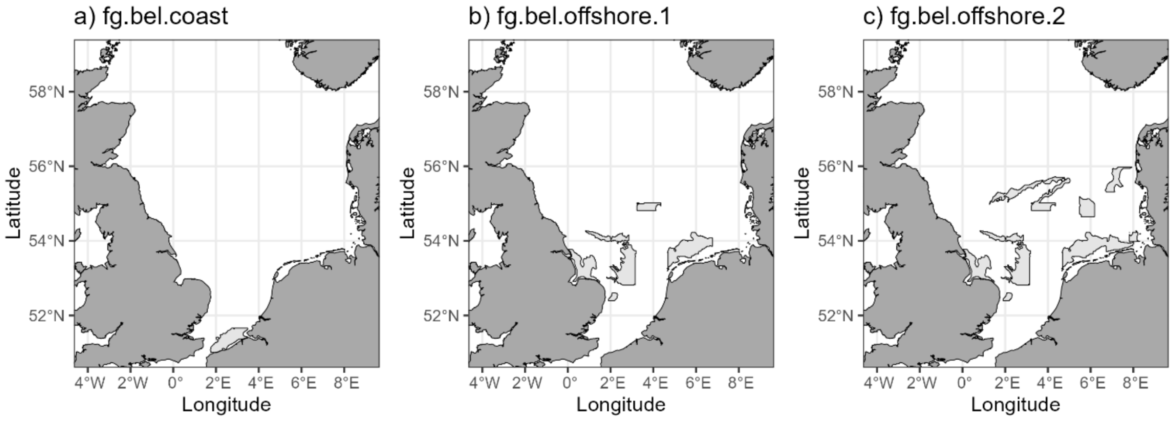

*Figure 5.3. Belgium. Fishing grounds of the bottom trawling fleets*

#### Denmark

Trawling grounds of Danish anchor seiners are shown in Figure 5.4 (***fg.fleet.dk.gpkg***). Grounds were obtained by digitizing the polygons directly from the published maps (Drechsel, 1890; Ministry of Agriculture, 1897-1915; Mortensen and Strubberg, 1935). Trawling grounds expanded from the norther coast of Jutland Figure 5.4a toward the west coast Figure 5.4b. In the first decade of the 20$^{th}$ century, anchor seining was concentrated in the core fishing ground *fg.dk.1900.core*, whereas the remaining effort took place in the peripheral area *fg.dk.1900.peri* (Figure 5.4c).

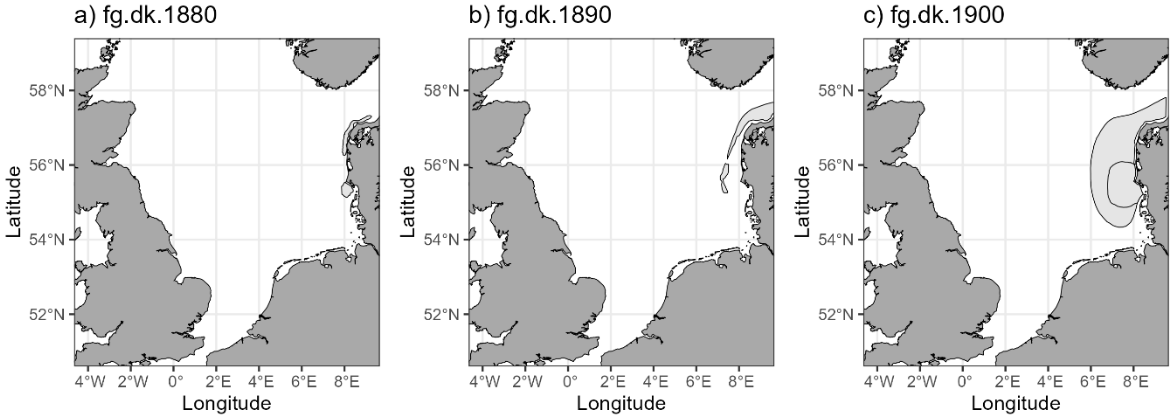

*Figure 5.4. Denmark. Fishing grounds of the anchor seining fleets in the 1880s, 1890s and 1900s. The fishing ground of the 1900s (fg.dk.1900) comprise of a core (dark) area with 50% of the effort (fg.dk.core) and a peripheral (light) area (fg.dk.peri).*

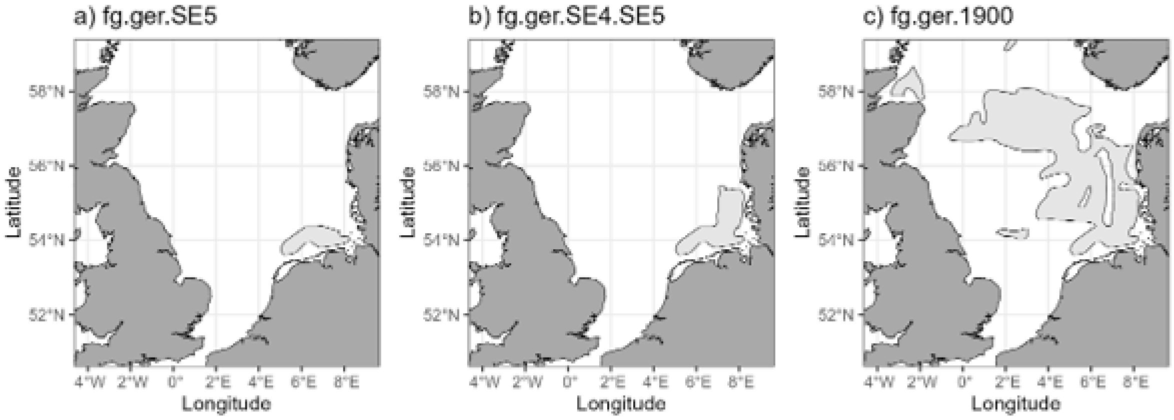

*Figure 5.5. Germany: fishing grounds of sail trawlers and steam trawlers in different time periods*

#### Germany

Fishing grounds of German fisheries were reported by Schnakenbeck (1927). Trawling grounds German bottom trawlers are shown in Figure 5.5 (***fg.fleet.ger.gpkg***). The trawling season of German sail trawlers started in March targeting plaice on the coastal grounds off Terschelling and Ameland. As the season progressed, trawling shifted eastwards, targeting sole, turbot, plaice and lobster (Darmer, 1894; Schnakenbeck, 1927) (Figure 5.5a). At the end of the century the fishing grounds included the north Frisian islands and the fishing season extended to November (Figure 5.5b). North Sea fishing grounds of steam trawlers were located in offshore waters of the central and northern North Sea (Figure 5.5c).

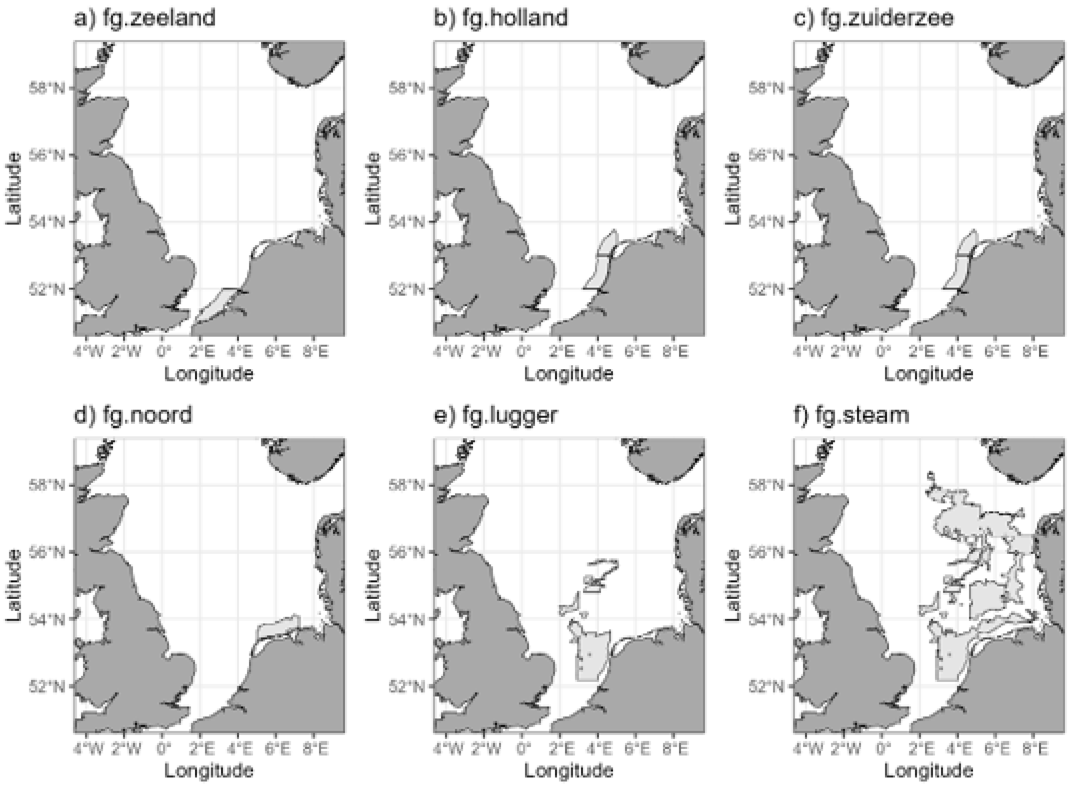

*Figure 5.6. Netherlands: fishing grounds of the different fleets of sail and steam bottom trawlers*

#### Netherlands

***SM1.tables.xls/fg.deveen*** provides the depth range (fathom) and the geographic boundaries (degrees.minutes) of the individual grounds of Dutch motor and steam trawlers in de Veen (2024) and information on the fishing grounds of sailing fleets in the annual fisheries reports (Collegie voor de Zeevisscherijen, 1857–1911) and reports on specific fisheries (Venema, 1869; Schouten, 1942; Petrejus, 1954; Dorleijn, 1982; de Vink, 2012). Polygons of individual grounds are given in ***fg.veen.gpkg***.

Fishing grounds of the Dutch fleets are shown in Figure 5.6 (***fg.fleet.nl.gpkg***). Sail trawl fleets restricted their trawling activities to grounds within 25 nm from the coast. Vessels from Zeeland were mainly fishing off Zeeland (Figure 5.6a), although fishers from Arnemuiden occasionally trawled off Texel – Ameland in summer (Venema, 1869). The fishing grounds of the bomschuiten (Zijde: Ter Heide, Scheveningen, Katwijk, Noordwijk, Zandvoort, Wijk aan Zee, Egmond, Petten) were located in the immediate vicinity of the coast between the mouth of the Meuse and Texel (exceptionally as far as Borkum) (Figure 5.6b). The trawling depth ranged 10-20 fathom in spring. Later in the season until the start of the herring season, vessels trawled at 10-15 fathom, occasionally at 6-7 fathom (Petrejus, 1954; de Vink, 2012). The fishing grounds of the vessels from the Zuiderzee resembled those of the fishers from de Zijde (Figure 5.6c). Fishers from the northern provinces of Friesland (Ameland, Wierum, Moddergat) and Groningen (Zoutkamp) trawled in the coastal waters off the west Frisian islands (Texel – Borkum) (Figure 5.6d), occasionally as far east as the mouth of the Elbe and trawled generally closer to the coast than English smacks in that area (Venema, 1869). Luggers trawled both coastal and offshore grounds, targeting species such as sole, plaice and haddock (Figure 5.6e). Around the turn of the century, luggers restricted their trawling activities to the coastal grounds due to declining catches of plaice and haddock on the Dogger

(Collegie voor de Zeevisscherijen, 1902). Steam trawlers mainly trawled grounds north of IJmuiden (Schouten, 1942; de Veen, 2024) (Figure 5.6f).

Table 5.2. Surface area ($10^3$ km$^2$ and %) of rough ground by depth zone (0-100m, 0-200m) as mapped by Olsen (1883), Close (1920) and WMR (unpublished).

| Type of ground | Surface area (0-100 m) | | Surface area (0-200 m) | |
|---|---|---|---|---|
| | $10^3$ km$^2$ | % | $10^3$ km$^2$ | % |
| Olsen (1883) | | | | |
| Stones Rocks | 36.7 | 8.1 | 37.0 | 7.7 |
| Moorlog | 2.5 | 0.5 | 2.5 | 0.5 |
| Oysters | 26.6 | 5.9 | 26.7 | 5.6 |
| Mud ooze | 57.4 | 12.7 | 68.7 | 14.3 |
| Clay | 17.6 | 3.9 | 26.3 | 5.5 |
| Close (1920) | | | | |
| Very rough | 2.4 | 0.5% | 2.4 | 0.5% |
| Rough | 12.9 | 2.9% | 13.1 | 2.7% |
| Medium | 8.3 | 1.8% | 8.3 | 1.7% |
| Fair | 8.1 | 1.8% | 8.4 | 1.7% |
| Total | 31.7 | 7.0% | 32.2 | 6.7% |
| WMR (unpublished) | | | | |
| Reef areas | 43.4 | 9.6% | 44.2 | 9.2% |

#### Foul grounds

Fishers may safely tow their bottom trawls in sandy areas but will avoid areas where there is a risk to damage or lose their gear. Information on the nature and location of rough grounds were described by (Olsen, 1878; Olsen, 1885). The sea floor map published in 1883 (Olsen, 1883) showed areas of stones

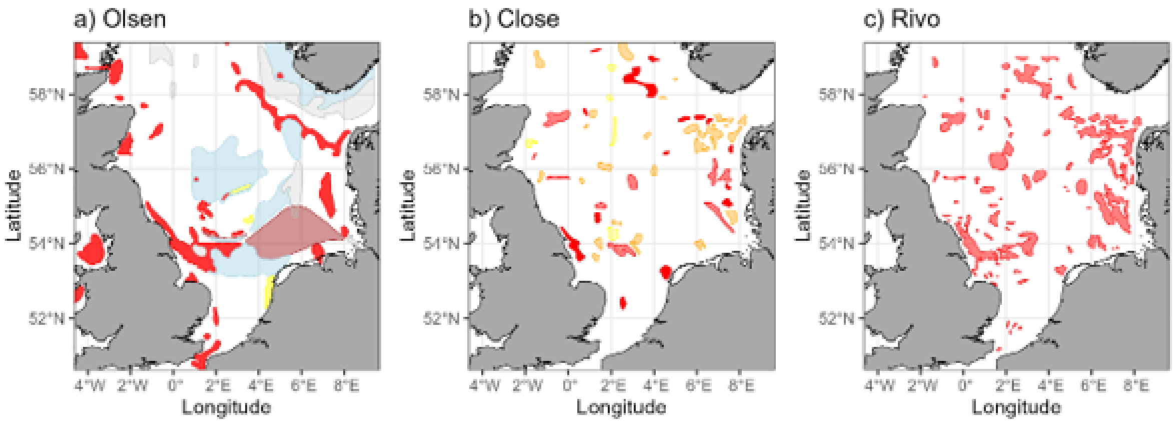

*Figure 5.7. Rough grounds according: a) Olsen (1883): stones and rocks (red), moorlog (yellow), mud ooze (light blue), oysters (brown), clay (light grey); b) Close (1920): very foul (red), foul (light red), medium (orange), fair (yellow); c) WMR (unpublished): reef areas.*

and rocks, moorlog, oysters, mud ooze, and clay as were known at that time (Figure 5.7a). Later maps from (Close, 1920; Figure 5.7b) and Netherlands Institute for Fisheries Research (WMR unpublished; Figure 5.7c) show, in general, similar patterns.

### 6. Swept area estimation by fleet

The total surface area of the seafloor swept annually by bottom trawlers SA is estimated as the sum of the swept areas of the different fleets $SA = \sum_f SA_f$.

$SA_f$ is a function of the number of vessels ($N_f$), the proportion of vessels engaged in bottom trawling ($P_f$), the maximum number of days at sea per year (das) of a vessel ($E_f$), the width of the gear ($G_f$), the towing speed ($U_f$), the number of trawling hours per day at sea ($H_f$) and the proportion of days with favourable conditions ($F_f$).

The input parameters used in the estimation of the swept area of the sail trawlers are given in separate worksheets in ***SM2.tables.xls***.

**tide.sediment.fleet** presents the mean tidal speed in the fishing ground of a fleet. This parameter was used to estimate the proportion of days with favourable conditions ($F_f$).

**sail.coefficients** presents the sail coefficients $Cx$ and $Cy$ used to estimate the aerodynamic thrust force and side force.

**n.vessel.sail** and **n.vessel.steam** presents the number of sail and steam trawlers by year and fleet ($N_f$).

**parm.sail.fleet** and **parm.steam.fleet** presents the following operational parameters of the sail and steam trawlers by fleet and time period:

| Code | Parameter | parm.sail | parm.steam |
|---|---|---|---|
| Country | country of origin | x | x |
| fleet.ID | fleet identifier | x | x |
| mode | mode of operation (single boater – fleeting; petite – grande) | x | |
| v.type | vessel type (smack, sloop, bomschuit) | x | |
| gear | gear type (beam trawl, otter trawl) | | x |
| start, end | time period | x | x |
| prop.nsea | proportion of the vessels that are trawling in the North Sea | x | x |
| prop.mode | proportion of vessels in mode-class | x | |
| prop.gear | proportion of vessels in gear-class | | x |
| trip.days | duration of a fishing trip (days) | x | x |
| trip.days.harbour | time spend in harbour between successive trips | x | x |
| month.year | number of month that the fleet is active during a year | x | x |
| das.year | number of days at sea (das) per year | *(1)* | x |
| sd.das.year | standard deviation of das.year | | x |
| towspeed.knot | towing speed (knot) | x | x |
| fish.hr.day | number of fishing hours per day at sea | x | x |

| | | | |
|---|---|---|---|
| loa.m | vessel length (length overall, m) | x | |
| tonnage | vessel size (ton) | x | |
| beam.m | width of the hull (m) | x | |
| n.trawl | number of beam trawls | x | |
| gear.width.m | Gear width (horizontal net opening). | *(2)* | x |
| fg.ID | fishing ground identifier | x | x |

*(1) The maximum number of days that a sailing vessel can be out at sea was set at 280 days (240-320) for all fleets; (2).Gear width for sail trawlers was estimated from the isometric relationship with vessel size (loa).*

If during a time periods a change occurred in an input parameter - for instance if the mode of operation changed from single boating to fleeting, or if the gear changed from beam trawl to otter trawl, or if the proportion of the fleet operating in the North Sea changed - a linear change in the parameter was assumed over the duration of the period.

**parm.anchorseine** presents the parameters used to estimate the swept area of the Danish anchor seiner fleet. $SA = N_{days}N_{hauls}S$ , where $N_{days}$= number of fishing days, $N_{hauls}$ = number of hauls per day, $S$ = the surface area swept in a single haul. The number of fishing days for the period 1897 to 1914 data originate from registered number of fishing day for Esbjerg vessels landing their catch in Esbjerg (Ministry of , 1897–1915); for the period 1876 to 1896 data is a mean of number of fishing days in the years 1897 to 1908. It is unclear why the number of days in 1898, 1899 and 1900 is about twice as low as in years before and after.

The number of hauls per day at sea is set at 8 hauls for the day light period (Blegvad, c1950).

The area swept by an anchor seine (S) is determined by the rope length (L) and was estimated following (Eigaard et al., 2016). Assuming an isosceles shape of the anchored seine, the height (h) of the isosceles triangle (gear footprint) and the swept area of the gear footprint (S) is given by

$$h = \sqrt{\left(\frac{L}{2}\right)^2 - \frac{\left(\frac{L}{4}\right)^2}{4}}$$

$$S = \frac{1}{2}\left(\frac{L}{4}\right)h$$

Rope length was estimated from a weighted least-square based on median the rope length from literature documenting minimum and maximum rope length in the years 1888 and 1929 (Mortensen and Strubberg, 1935). Rope length before 1888 was fixed to the median between the minimum and maximum documented rope length in 1888.

### 7. Sensitivity analysis

The sensitivity of the swept area estimate was explored by estimating the effect of a 10% increase or decrease in an input parameter on the swept area estimate for a selection of four sail trawling fleets (Figure 7.1). The simulation shows that a 10% change in vessel size has an equally large effect on the estimated swept area in all fleets (Figure 7.1c). In contrast, towing speed and beam trawl width have a different effect. For the Thames fleet, the swept area is almost unaffected by a change in towing speed, whereas in the bomschuit of the Zijde SKN fleet, a 10% change in towing speed had a larger and opposite effect on the estimated swept area. The Humber and German ewer fleet were intermediate (Figure 7.1a). A 10% change in beam trawl width results in a change in swept area in the same direction. The strength of the effect is slightly smaller in Thames and Zijde fleets (about 8%) and substantially smaller in the Humber (<=2%).

The difference in sensitivity between the fleets is related to the interaction between the direct effect of towing speed on the swept area and the higher wind force required to tow the gear at the higher speed. The latter will reduce the proportion of days with favourable winds (trawling opportunity window). The degree to which this will be reduced will depend on the vessel dimensions and strength of the tidal currents at the fishing grounds.

To further explore the sensitivity, we estimated the combined effect of towing speed, tidal speed and beam trawl width on the swept area after taking account of the effect on the proportion of favourable days (Figure 7.2). The effect was expressed relative to the swept area at the typical towing speed (2 knots), tidal seed (1 knot) and gear width (12m) of a sailing smack of 19.5m (loa). The reference is illustrated by the white dot in panel B of Figure 7.2. The analysis showed that an increase in towing speed reduced the swept area, in particular at lower levels of tidal speed, due to a reduction in the proportion of favourable days. The negative effect on the proportion of favourable days increase with the size of the beam trawl.

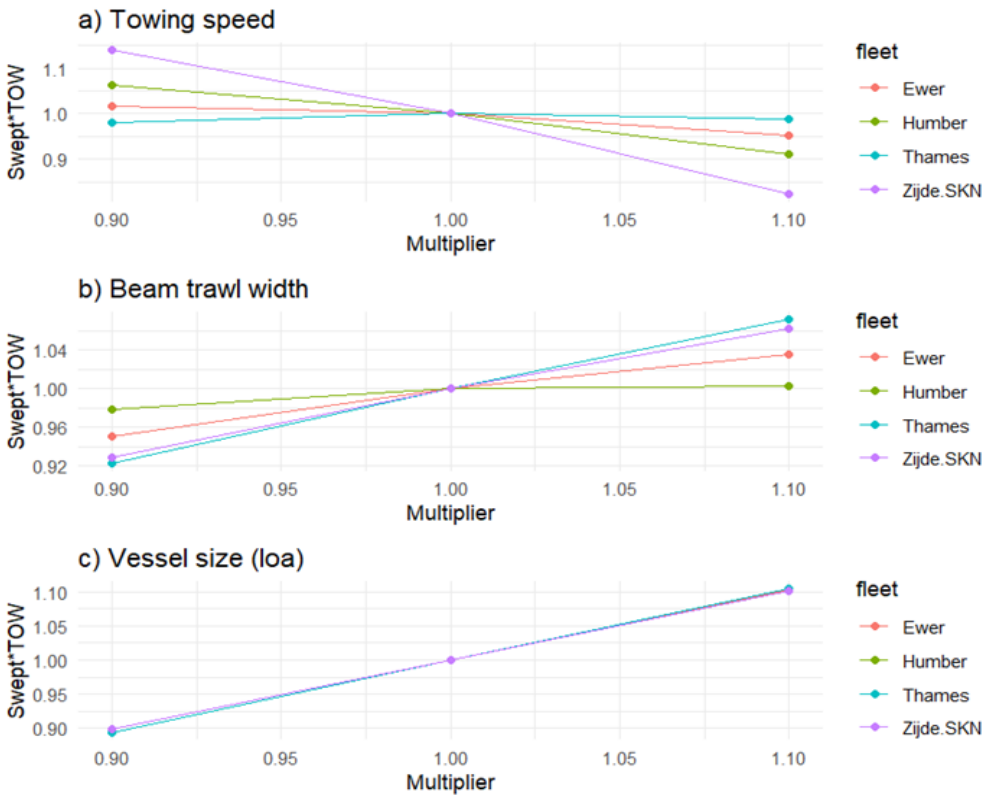

*Figure 7.1 Sensitivity analysis. Effect of a 10% change in input parameters (Towing speed, Beam trawl width (BTL), Vessel size) on the swept area estimate for a selection of four sail trawling fleets. Swept area (Swept.TOW) was estimated taking account of the differences in towing speed, gear width and their effect of the proportion of favourabe days (trawling opportunity window). corrected for the proportion of favourable days.*

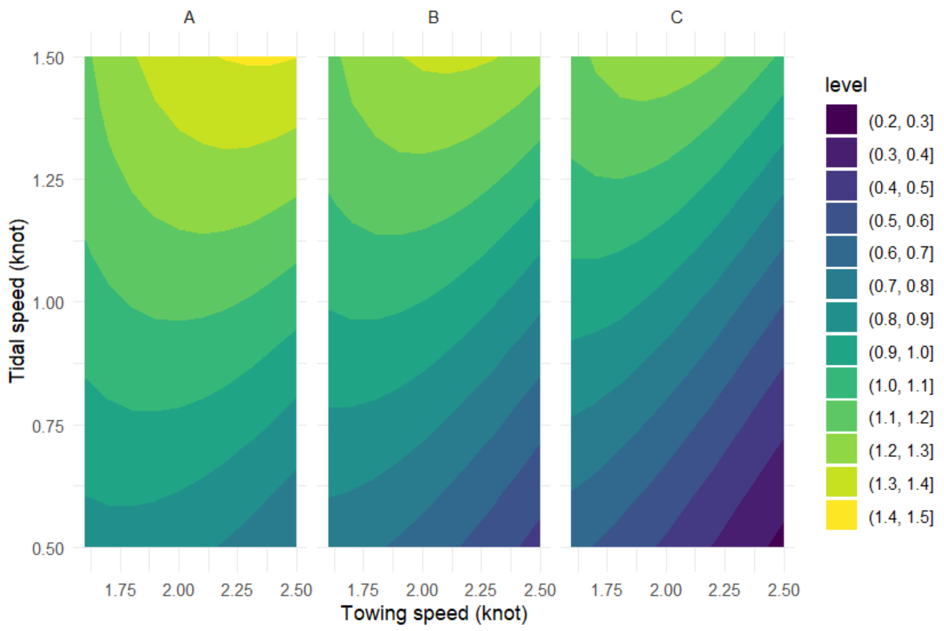

*Figure 7.2 The effect of combinations of towing speed (x-axis), tidal speed (y-axis) and beam trawl length (BTL=9m - panel A, BTL=12m – panel B and BTL=15m – panel C) on the swept area of a 19.5 m sailing smack. The effect ratio (level) was expressed relative to the swept area at towing speed = 2 knot, tidal speed = 1 knot and BTL = 12 m.*

### 8. Data tables

Data can be found in the following files at **10.5281/zenodo.22078260:**

- SM1.tables.xlsx
- SM2.tables.xlsx
- fg.olsen.gpkg
- fg.veen.gpkg
- fg.fleet.nl.gpkg
- fg.fleet.bel.gpkg
- fg.fleet.dk.gpkg
- fg.fleet.eng.sco.gpkg
- fg.fleet.ger.gpkg

Collegie voor de Zeevisscherijen (1854) Verslag over de zeevisscherijen. Van Weelden en Mingelen, 's-Gravenhage. 405p.

Collegie voor de Zeevisscherijen (1857-1911) Verslag (omtrent / van) den staat der zeevisscherijen. Van Weelden en Mingelen, 's-Gravenhage.

Collins JW (1889) Beam-trawl fishery of great Britain, with notes on beam=trawling in other European countries. Bulletin of the United States Fish Commission for 1887. Government Printing Office, Washington

Darmer JD (1894) Segel-handbuch fur Nordsee-fischer. Ein Kommentar zur Karte: Die Norsee Fischerei Grunde., Deutschen Seefischereiverein. 122 pp.

de Veen J (2024) De trawlvisserij en haar ontwikkeling. Een ongepubliceerd manuscript geschreven in de periode 1941-1945. Wageningen Marine Research, Wageningen Marine Research rapport 24.003. 114 pp.

de Vink L (2012) Aan boord van een Katwijkse bomschuit in de achttiende eeuw. Het handschrift van schipper Leendert Buijsertszoon van der Plas. Primavera Pers, Leiden.

de Zuttere C (1909) Enquete sur la peche maritime en Belgique: introduction, recencement de la peche maritime. , Lebegue & cie, Bruxelles. 634 pp.

Dorleijn P (1982) Van gaand en staand want. De zeilvisserij voor en na de afsluiting van de Zuiderzee. I. van Kampen & Zn., Bussum

Drechsel CF (1890) Oversigt over vore Saltvandsfiskerier I Nordsøen og Farvandene indenfor Skagen med Kort og Planer. Dansk Fiskerimuseum.

Eigaard OR, Bastardie F, Breen M, Dinesen GE, Hintzen NT, Laffargue P, Mortensen LO, et al. (2016) Estimating seabed pressure from demersal trawls, seines, and dredges based on gear design and dimensions. ICES Journal of Marine Science, 73: i27–i43.

Fishery Board for Scotland (1894) Twelfth Annual Report of the Fishery Board for Scotland, being for the year 1893. Part III. Scientific investigations. 410 pp.

Garstang W (1900) The impoverishment of the sea - a critical summary of the experimental and statistical evidence bearing upon the alleged depletion of trawling grounds. J Mar Biol Assoc UK 6:1–69

Holdsworth EWH (1874) Deep-sea fishing and fishing boats. Edwards Stanford, London, England

ICES (1956) Bulletin statistique des pêches maritimes, Copenhagen.

Kramer R (1984) De visserij van de zijde in de eerste helft van de 19e eeuw. Holland Regionaal-historisch tijdschrift 16:175–188

le Compte L (1831) Afbeeldingen van schepen en vaartuigen. F. Kaal, Amsterdam

Lescrauwaet A-K, Fockedey N, Debergh H, Vincx M, Mees J (2013) Hundred and eighty years of fleet dynamics in the Belgian sea fisheries. Rev Fish Biol Fish 23:229–243

March EJ (1953) Sailing trawlers. The story of deep-sea fishing with long line and trawl. Percival Marshall and Company Ltd, London.

March EJ (1970) Sailing Trawlers. The Story of Deep-Sea Fishing with Long Line and Trawl, 2nd edition. Newton Abbot, David & Charles.

McLaverty C, Roberts CM, Bennema FP, Hunt GL, Maynard E, Moore ABM, Phillips E, et al. (2026) The industrialisation of the seabed: Estimating century-scale impacts of bottom trawling on shelf sea sediments. Preprint from Research Square, https://doi.org/10.21203/rs.3.rs-9640267/v1

Masterman AT (1914) Report on the plaice fisheries of the North Sea (Parts I and II). Fishery Investigations, Series II, Vol II:120

Ministry of Agriculture (1897-1915). Annual fisheries reports covering the years 1895-1914. Copenhagen: G. E. C. Gad.

Mortensen FV and Strubberg AC (1935) Dansk Saltvandsfiskeri. C.A. Reitzel. Copenhagen.

Olsen OT (1878) The fisherman's practical navigator. Walker & Brown, Highstreet, Hull., Hull.

Olsen OT (1883) The piscatorial atlas of the North Sea, English and St. George's Channels, illustrating the fishing ports, boats, gear, species of fish (how, where, and when caught), and other information concerning fish and fisheries. Taylor and Francis, London 58p..

Olsen OT (1885) The fisherman's practical navigator. Second Edition: illustrated, revised, and improved., O.T. Olsen, Grimsby. J. Imray & Son, Minories, London. 362 pp.

Orban de Xivry J (1892) Etude sur la grande pêche maritime Belge. Imprimerie Ch. Lemaire: Liège. XII, 273 pp.

Petrejus EW (1954) De bomschuit, een verdwenen scheepstype. Publicaties van het Museum van Land- en Volkenkunde en het Maritiem Museum "Prins Hendrik" Book 1954. No 2. M.M.1, Rotterdam

Redeke HC (1907) Rapport over onderzoekingen betreffende de visscherij in de Zuiderzee ingesteld in de jaren 1905 en 1906. Ministerie van Landbouw, Nijverheid en Handel., 's-Gravenhage

Robinson R (2000) The line and trawl fisheries in the age of sail. In: Starkey DJ, Reid C, Ahscroft N (eds) England's Sea Fisheries. Chatham Publishing, London

Royal Commission (1866) Report of the Commissioners appointed to inquire Into the sea fisheries of the United Kingdom. Volume I & II. 1590 pp. Eyre and Spottiswood, London.

Royal Commission (1885) Trawl net and beam trawl fishing. Report of the commissioners appointed to inquire and report upon the complaints that have been made by line and drift net fishermen of injuries sustained by them in their calling owing to the use of the trawl net and beam trawl, in the territorial waters of the United Kingdom; with minutes of evidence and appendix. 564 p. Eyre and Spottiswood, London.

Royal Commission (1893) Report from the Select Committee on Sea Fisheries; Together With the Proceedings of the Committee, Minutes of Evidence, Appendix And Index. Eyere and Spottiswoode, London. 496 p.

Schnakenbeck W (1927) Die Nordseefischerei. Handbuch der Seefischerei Nordeuropas Band V Heft 1. Schweizerbart, Stuttgart

Schouten A (1942) De Nederlandsche Groote Trawlvisscherij. PhD Utrecht, Rijksuniversiteit Utrecht, Utrecht

Select Committee (1833) Special Report from the Select Committee on the British Channel Fisheries with Minutes of Evidence and Appendix. The House of Commons, London. 168 p.

Select Committee (1900) Special Report and Report From the Select Committee on the Sea Fisheries Bill; Together With the Proceedings of the Committee, Minutes of Evidence, Appendix and Index. Wyman and Sons, London. 196 p.

Timmermann G (1962) Die nordeuropäischen Seefischereifahrzeuge, ihre Entwicklung und ihre Typen. Handbuch der Seefischerei Nordeuropas Band XI, Heft 4. Schweizerbart, Stuttgart

UK110. Zeilplan en lijnentekening van de schokker “ UK 110” gebouwd in 1883 door Fledderus te Kuinre. https://boekenplank.ssrp.nl/beschrijving-van-de-oude-scheepstypes/kenmerk-beschrijvingen-schokkers-bonzen-en-pluten. Accessed 23 August 2026.

Venema GA (1869) De visscherij in de provincie Groningen. In: Bijdragen tot de kennis van den tegenwoordigen staat der provincie Groningen. Gebr. Hoitsema, Groningen: 83-271.

Verbrugghe J (1932) Die Belgische Seefischerei. In Handbuch der Seefischerei Nordeuropas. Band VII, Heft 3. Schweizerbart, Stuttgart.

Versteeg WK (1947) Scheepsmodellen 1700-1900: bevattende modellen van visschersschepen, vrachtschepen, schepen voor groote vaart, yachten, enz. Nederlandsche boekhandel. Antwerpen.

**Part 2. 10.5281/zenodo.21904478**

### The mechanics of sail trawling

Adriaan D. Rijnsdorp[1)]; Floris P. Bennema[2)]; Frans Veenstra[3)]; Ole R. Eigaard[4)]; Jasmin Ann-Christine Thomassen[4)]; Ciarán McLaverty[5)]

[1)]Wageningen Marine Research, PO Box 68, 1970 AB IJmuiden, The Netherlands; [2)]MarHis, Haren, The Netherlands; [3)]VFC Veenstra fishery consultancies, Haarlem, The Netherlands; [4)]National Institute of Aquatic Resources (DTU Aqua), Technical University of Denmark, DK-2800 Kongens Lyngby, Denmark; [5)]Centre for Ecology and Conservation, University of Exeter, Penryn Campus, Cornwall, TR10 9FE, England.

#### Abstract

Bottom trawls have been used for centuries, yet studies of their impact on marine ecosystems have largely been restricted to recent decades. Bottom trawling before the advent of steam and diesel engines was dependent on wind power. In order to quantify bottom trawling effort of historic sailing vessels we need to know the minimum wind speed required to tow the fishing gear over the sea floor. Here we derive an approach to estimate the minimum wind requirements based on information on the characteristics of the sailing vessel and the fishing gear. The method is used in a study that reconstruct the fishing effort of the international bottom trawl fleets in the North Sea between 1820 and 1914 encompassing the age of sail to early steam trawling. The results are published in "From sailing to steam trawling: the evolution of bottom trawl effort in the North Sea".

#### 1. Aerodynamic thrust and side force

The mechanics of sailing is a complex interplay of hydrodynamic and aerodynamic forces. As we are interested in a first-order approximation to estimate the minimum wind force required for trawling that can be applied to different types of sailing vessel, we use a simplified mathematical representation of dominant processes.

The thrust force that will push a sailing vessel forward is generated by the force of the wind on the sails. The wind force on the vessel is combined effect of the true wind and the wind generated by the movement of the vessel (sail wind). If $V_t$ = true wind, $V_b$ = vessel speed, θ = angle between the true wind and the course of the vessel, the apparent wind $V_a$ is given by,

$$V_a = \sqrt{{V_t}^2 + {V_b}^2 + 2V_tV_bcos(\theta)} \qquad [1.1]$$

Figure 1 illustrates the equilibrium between the aerodynamic ($F_A$) and hydrodynamic ($F_H$) forces in the horizontal plane for a vessel sailing at a constant speed (Slooff 2015). In equilibrium, thrust force $T$ is equal to, but in opposite direction, to the hydrodynamic resistance $R$, whereas the aerodynamic side force $S_A$ is equal to, but in opposite direction to the hydrodynamic side force $S_H$.

If $V_A$ = apparent wind speed, $A$ = sail surface area, $\rho_{air}$ = density of the air, and $C_X$ and $C_Y$ are the aerodynamic coefficients, the thrust force $T$ and the side force $S_A$ are given by

$$T = \frac{1}{2}\, C_X\, A\ \ \rho_{air}\, {V_a}^2 \qquad [1.2]$$

$$S_A = \frac{1}{2}\, C_Y\, A\ \ \rho_{air}\, {V_a}^2 \qquad [1.3]$$

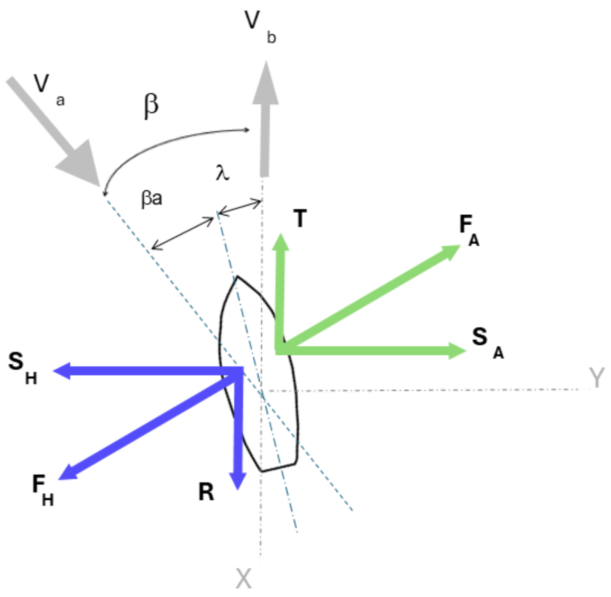

*Figure 1. Equilibrium of aerodynamic ($F_A$) and hydrodynamic ($F_H$) forces in the horizontal plane of a sailing vessel at a constant speed. $V_a$ and $V_b$ are the vectors of the apparent wind speed and boat speed. T and $S_A$ are the vectors of the aerodynamic thrust force and side force. R and $S_H$ are the vectors of the hydrodynamic resistance and side force. $\beta$ is the angle between the apparent wind and the course of the vessel. $\beta_a$ is the angle between the apparent wind and the vessel axis. $\lambda$ is the leeway angle between the vessel axis and the course of the vessel which is mainly due to the moment generated by the side forces. The coordinate system is aligned with the vessel course (X axis). Equilibrium of moments is not shown. Adapted from Slooff (2015).*

The aerodynamic coefficients $C_X$ and $C_Y$ are functions of the angle β between the apparent wind and the course of the vessel. The relationships differ between sail types (main sail, jib, etc). We used an overall relationship estimated for a broad range of sail types (Figure 2; Anon 2025).

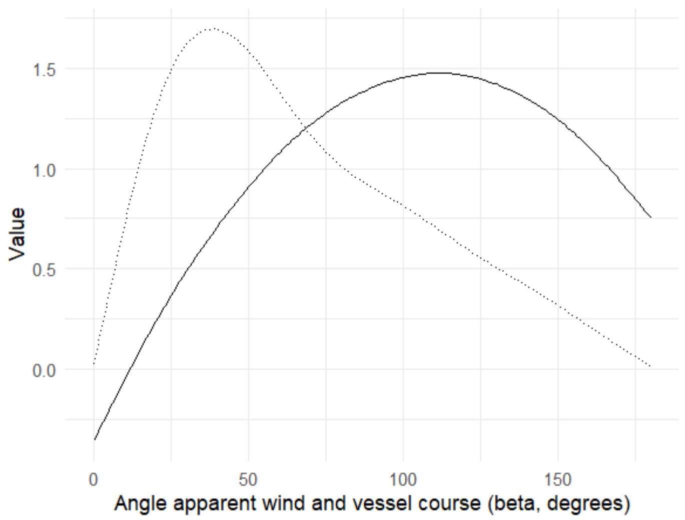

*Figure 2. Coefficients of the thrust force (CX full line) and lateral force (CY dotted line) in relation to the angle $\beta$ between the apparent wind and the vessel course.*

### 2. Hydrodynamic resistance

The hydrodynamic resistance of a sailing vessel ($R_{total}$) comprises several components that are related to different mechanisms.

$$R_{total} = R_i + R_F(1 + k_1) + R_A + R_W \qquad [2.1]$$

The $R_i$ component represents the resistance induced by pressure difference on the hull due to the aerodynamic side force $S_H$ and is specific for sailing vessels (Slooff 2015). The other components, which apply to sailing and motorized vessels, are based on (Holtrop & Mennen 1982).

$R_i$ is approximated by

$$R_i = \frac{S_A^2}{0.5\,\rho_W\,{V_b}^2\,e\,\pi\,D^2} \qquad [2.2]$$

Where $\rho$ = density of the water, $V_b$ = vessel speed in water, $D$ = mean draught of the hull, $e$ = efficiency of the hull to produce the hydrodynamic side force (Slooff 2015, Larsson et al. 2022).

The frictional resistance drag ($R_F$) is given by

$$R_F = \frac{1}{2}\rho C_f A {V_b}^2 \qquad [2.3]$$

where $A$ = wetted surface area of the hull, $V_b$ = boat speed relative to the water. The friction coefficient $C_f$ is calculated based on the Reynolds number $Re$ using the ITTC-1957 relation

$$C_f = 0.075/(log Re - 2)^2 \qquad [2.3.1]$$

The form factor $(1 + k_1)$, applicable for vessels with a T/L ratio >0.05 and a distance *lcb* = -1.5% between the centre of buoyancy forward of 0.5L as a percentage of L, is given by

$$1 + k_1 = 0.93 + \left({}^{T}/_{L}\right)^{0.2228446}\left({}^{B}/_{L_R}\right)^{0.92497}(0.95 - C_P)^{-0.521448}(1 - C_P - 0.0357143)^{0.6906} \qquad [2.3.2]$$

where $T$ = mean draught, $L$ = length of the water line, $B$ = beam of the water line, $C_P$ = prismatic coefficient and $L_R$

$$L_R = L\left(1 - C_P - 0.095\,{}^{C_P}/_{(4C_P - 1)}\right) \qquad [2.3.3]$$

The model-ship correlation resistance ($R_A$) with coefficient $C_A$ corrects for the roughness of the hull and the still-air resistance for roughness values higher than $k_s = 150\mu m$.

$$R_A = \frac{1}{2}\rho {V_b}^2 S C_A \qquad [2.3.4]$$

with $C_A = \left(0.105{k_s}^{1/3} - 0.005579\right)/L^{1/3}$ [2.3.5]

The wave-making resistance ($R_w$) is a function of displaced volume of water ($\nabla$), water density ($\rho$), gravitational constant (g), and the Froude number ($F_n$), based on the length of the water line ($L$) and the boat speed relative to the water ($V_b$) and coefficients $c_1, m_1, m_2$ described below:

$$R_w = c_1 \nabla \rho g\, exp(m_1 F_n^{-0.9} + m_2 \cos(\lambda F_n^{-2}) \qquad [2.4]$$

$$F_n = {}^{V_b}\Big/_{\sqrt{gL}} \qquad [2.4.1]$$

$\lambda = 1.446\ C_p - 0.03\ {}^{L}/_{B}$ (when L/B < 12)
[2.4.2]

Coefficient $c_1$ is given by

$$c_1 = 2223105\ {c_7}^{3.78613} \left({}^{T}/_{B}\right)^{1.07961} (90 - i_E)^{-1.27565} \qquad [2.4.3]$$

with

$$c_7 = 0.5 - 0.0625\ {}^{L}/_{B} \qquad [2.4.4]$$

Coefficient $i_E$ is the angle of the waterline at the bow in degrees. It can be calculated from the equation below which is based on a regression analysis of 200 hull shapes.

$$i_E = 1 + 89 \exp\left\{-{}^{L}/_{B}{}^{0.80856} (1 - C_W)^{0.30484} (1 - C_P - 0.0225 lcb)^{0.34574} \left({}^{100\nabla}/_{L^3}\right)^{0.16302}\right\} \qquad [2.4.5]$$

Coefficient $m_1$ is given by

$m_1 = 0.0140407\ {}^{L}/_{T} - {}^{1.75254\nabla^{1/3}}/_{L} - 1.79323\ {}^{B}/_{L} - c_{16}$
[2.4.6]

with

$$c_{16} = 8.07981 C_p - 13.8673 {C_p}^2 + 6.984388 {C_p}^3 \quad \text{(when } C_p<0.80\text{) or} \qquad [2.4.7]$$

$$c_{16} = 1.73014 - 0.7067 C_p \quad \text{(when } C_p>0.80\text{)} \qquad [2.4.8]$$

Coefficient $m_2$ is given by

$$m_2 = -1.69385\ C_p^2\ e^{-0.1 F_n^{-2}} \qquad [2.4.9]$$

### 3. Wind speed threshold

To estimate the wind speed threshold required for a sailing vessel to tow a beam trawl at a constant speed the thrust force $T$ (equation [1.2]) should be in equilibrium with $R_i$, the resistance induced by the aerodynamic side force (equation [2.2]), and $G$, the sum of the drag of the beam trawl towed at the vessel speed ($V_b$) and other resistance components

$$T = R_i + G \qquad [3.1]$$

The apparent wind speed $V_a$ required to tow the beam trawl at a given $V_b$ is given by equation [3.2] that is obtained by substituting eq[1.2] and eq[2.2] in eq[3.1] and rearranging for $V_a$

$$V_a^2 = \frac{b c_x \pm \sqrt{b^2 {c_x}^2 - 4\, {c_y}^2\, b\, G}}{2\, {c_y}^2} \qquad [3.2]$$

where $c_x = \frac{1}{2} C_X\, A\, \rho_{air}$; $c_y = \frac{1}{2} C_Y\, A\, \rho_{air}$ and $b = 0.5\, \rho_{water}\, {V_b}^2\, e\, \pi\, D^2$

The true wind speed was calculated as

$$V_t = \sqrt{V_a^2 + V_b^2 - 2 V_a V_b \cos\beta} \qquad [3.3]$$

where $\beta$ is the apparent wind angle.

The true wind angle is calculated as

$$\theta = \text{atan}(V_a \sin(\beta)/(V_a \cos(\beta) - V_b)) \quad [3.4]$$

### 4. Gear drag

#### 4.1. Hydrodynamic drag of gear components

The hydrodynamic drag of a beam trawl can be estimated by summing the drag of the gear components following (Paschen et al. 2000, O'Neill & Ivanović 2016, O'Neill & Summerbell 2016, Rijnsdorp et al. 2021). In brief, the hydrodynamic drag of a gear component ($D$) is related to the frontal surface area of the component ($A_f$), the towing speed ($U$), the density of the fluid ($\rho$) and the drag coefficient ($C_D$) specific for the geometry of the gear component, the Reynolds number and the surface roughness.

$$D = 0.5\rho A_f U^2 C_D \quad [4.1]$$

When objects are towed at an angle to the flow, the component of the hydrodynamic pressure forces aligned with the velocity component in the towing direction is considered. For the ground rope, being towed freely from the shoes, the angle of attack differs between the various segments. By decomposing the ground rope into segments, the drag of each component was calculated and summed as an estimate of the drag of the ground rope.

The geometry of the ground rope can be described by a catenary with $x$ and $y$ coordinates described by

$$y = a \cos h\left(\frac{x}{a}\right) \quad [4.2]$$

where $a$ is a solution to

$$2\alpha a \sin h\frac{L}{2a} = S \quad [4.3]$$

where $L$ is the length of the ground rope and $S$ is the distance between the two hanging points that equals the width of the beam trawl. The drag of a cylindrical rope segment of length $l$, diameter $d$, and angle of attack $\alpha$ is given by

$$D_c = 0.5\rho U^2 dlC_d sin^3(\alpha) \quad [4.4]$$

The drag of the net panels was estimated using the empirical results of (Reid 1977) showing that the drag was dependent of the surface area of the twine of the net (A) and the towing speed (U)

$$D_n = \frac{0.5\rho U^2 A*0.643}{1+0.923U} \quad [4.5]$$

$$A = \frac{N_t+N_b}{2} N_d M d \quad [4.6]$$

where $N_t$ is the number of meshes on the top row of the panel, $N_b$ is the number of meshes of the bottom row, $N_d$ is the number of meshes along the length of the panel, $M$ is the stretched mesh size, and $d$ is the twine diameter (Ferro 1981).

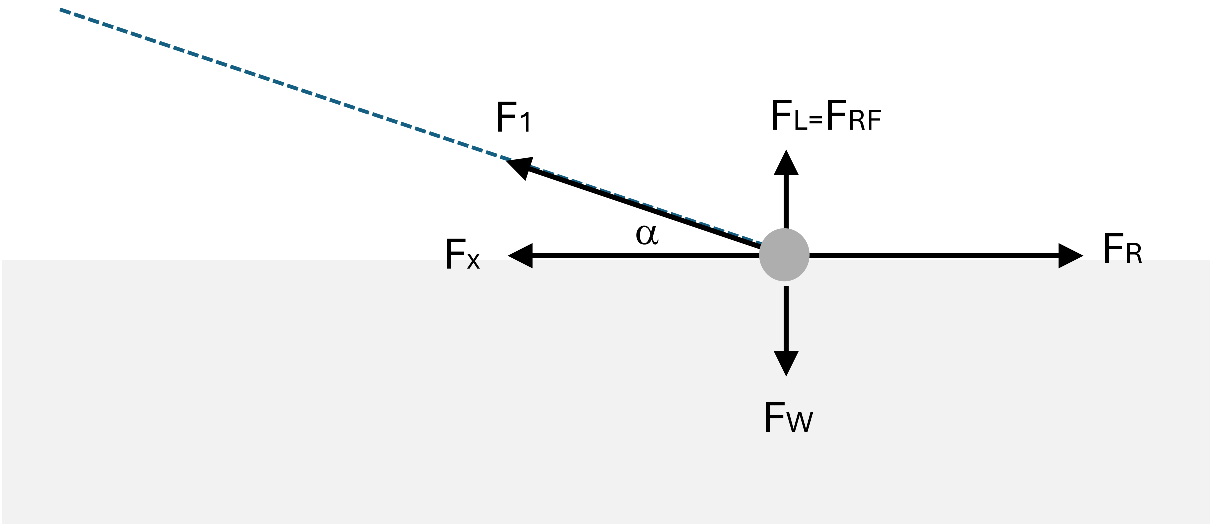

*Figure 3. Forces acting on a beam trawl that is towed over the sea floor. F1 shows the towing force of the trawl warp on the beam trawl. $F_x$ shows the towing force component in the direction of the movement. $F_L$ shows the upward component of the towing force. $F_R$ shows the total resistance force due to the geotechnical drag and the hydrodynamic drag of the beam trawl. $F_W$ is the force of the beam trawl on the sea floor due to the weight of the gear. $F_{RF}$ is the reaction force of the sea floor. The beam trawl is represented by the dark grey circle, the seafloor by the light grey rectangle and the trawl warp by the dashed line. dashed line. The dark grey circle represents the beam trawl. The dashed line shows the trawl warp. In our calculations we assumed that a fisher will adjust the gear, towing speed and warp to balance $F_W$ and $F_L$.*

#### 4.2. Geotechnical drag

The forces acting on a beam trawl towed over the sea floor is illustrated in Figure 3. F1 shows the towing force of the trawl warp on the beam trawl that can be decomposed in a component in the direction of the movement ($F_x$) and an upward component ($F_L$). $F_W$ is the normal force on the sea floor due to the weight of the gear and $F_{RF}$ is the reaction force of the sea floor. $F_R$ is the total resistance force due to the hydrodynamic drag of trawl components and the geotechnical drag.

The geotechnical drag ($F_{GT}$) can be described as the sum of the friction drag due to the pressure on the seabed ($F_w$), the passive pressure from the soil mass surrounding the gear ($F_{pp}$) and the dynamic force due to the acceleration of the soil mass in front of the gear ($F_d$) (Ghorai et al. 2025).

$$F_{GT} = K_1 F_w + K_2 F_{pp} + K_3 F_d \quad [4.7]$$

$$F_w = g\ M_{suspended} \quad [4.8]$$

$$F_{pp} = g\ (\rho_{soil} - \rho_{water})\ V \quad [4.9]$$

where $M_{suspended}$ is the mass of gear component in the water and $V$ is the volume of the sediment displaced by the gear component. (Ghorai et al. 2025) estimated the $F_{GT}$ of three configurations of tickler chain rigged beam trawls in the laboratory. They found that the dynamic force contribution was insignificant (K3 ~ 0). The other coefficients were estimated at K1 ~ 0.94 and K2 ~ 5.2. The predicted values were close to the measured values for the medium and heavy gear configuration but for the light gear configuration, the predicted values were about 1.5 – 2 too high. As the historic beam trawl gears were not rigged with tickler chains, we set coefficients K1 = 0.47 and K2=2.6 approximating the light gear.

##### 4.3. Total gear drag

The total drag ($F_R$) of a beam trawl is the sum of the hydrodynamic $F_{Hi}$ and geotechnical $F_{GTi}$ drag of the gear components *i*.

$$F_R = \sum_i F_{Hi} + \sum_i F_{GTi} \quad [4.10]$$

the effect of towing speed on the penetration of gear components into the sediment could not be incorporated in our model because of the complexity of these processes and the lack of a simple rule of thumb (Ivanović et al. 2011, O'Neill et al. 2018). We also ignored the effect of the friction drag (FD) component of the geotechnical drag, because we expect that the FD will be negligible because of the interaction between the weight of the gear component and the lift force and a fisher will adjust the warp length and towing speed to avoid the beam trawl to come lose from the seafloor.

#### 5. Gear and vessel resistance of a reference smack

This section presents the quantitative analysis of the aerodynamics and hydrodynamics of a smack that is representative for the English fleet that dominated the sail trawling in the 19$^{th}$ century (Table 1). A representative smack had a LOA = 20m, sail surface area = 200 m$^2$ and towed one 12m beam trawl at a speed of 2 knots over the sea floor, which corresponded to a towing speed in the water of 1 knot (range: 0.5 – 1.5 knot) (Holdsworth 1874, 1883).

The beam trawl used in the historic trawl fisheries is relatively light as compared to the present day beam trawl that has a median penetration depth of 2.7 cm (Hiddink et al. 2017). Assuming a penetration depth of 1 cm, the resistance of a beam trawl representative for the 1860s (Table 2) is estimated at 2.4 kN for a towing speed of 1 knot above the tide. Gear resistance is dominated by the hydrodynamic drag. The geotechnical drag gains importance at lower towing speed and higher penetration depth. The gear components with the largest contribution are the net and ground rope. The contribution of the warp, beam and shoes is relatively small. The contribution of the ground rope is highly dependent on the penetration depth as the shear force of the ground rope becomes increasingly important with an increase in penetration depth (Figure 4a). Gear resistance for a towing speed of 0.5 and 1.5 knot above the tide is estimated at 1.2 – 4.1 kN, respectively (Figure 4b).

Vessel resistance increased with the speed through the water and becomes increasingly dominated by the wave-making resistance (Figure 5a). At the typical towing speed through the water (between 0.5 – 1.5 knots), the contribution of the wave making resistance is negligible and resistance is mainly determined by the friction and viscous (form) drag component. The hydrodynamic resistance induced by the aerodynamic side force (Ri) becomes increasingly important when the vessel sails closer to the wind (Figure 5b).

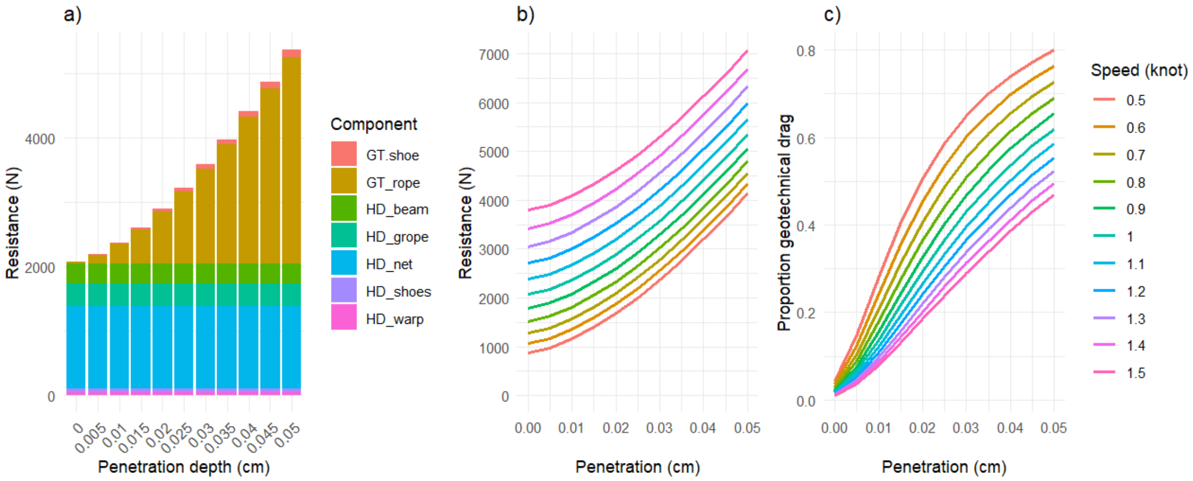

*Figure 4. a) contribution of the gear components to the total resistance of a 12 m beam trawl towed at 1 knot above the tidal (2 knot over the sea bed) in relation to penetration depth; b) effect of towing speed and penetration depth on the resistance force; c) relative proportion of the geotechnical resistance of the total resistance in relation to the penetration and towing speed.*

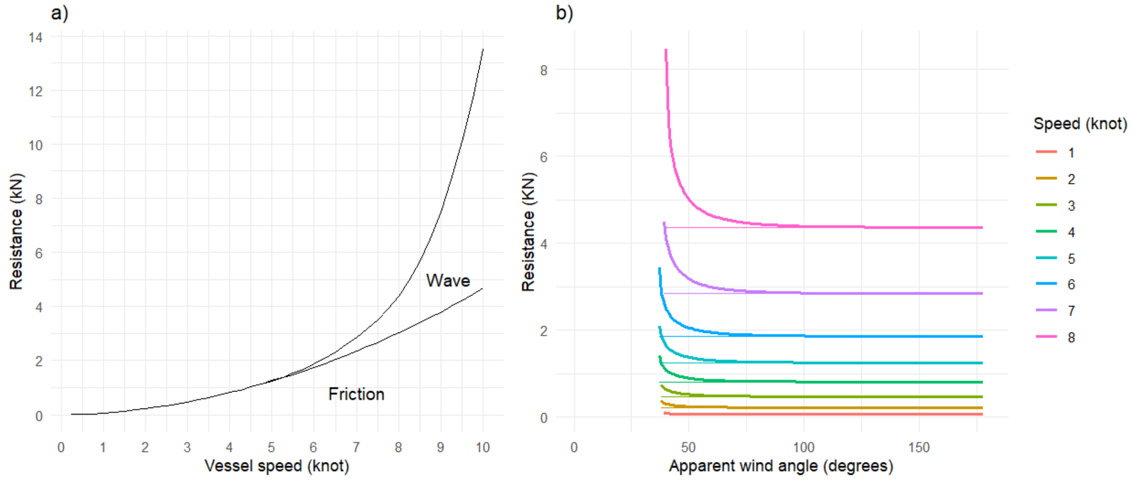

*Figure 5. a) Contribution of the friction drag and wave induced drag to the vessel resistance (kN) of the reference sail trawler at different sailing speeds; b) Vessel resistance (heavy line) and resistance due to friction and wave induced drag (thin line) of the reference trawler as a function of the angle between the apparent wind and the vessel course. The difference between heavy and thin line shows the contribution of the hydrodynamic resistance induced by the aerodynamic side force (Ri).*

The results in Figure 5 can be used to infer the gear resistance from anecdotal information of the reduction in sailing speed after launching a beam trawl. Smacks fishers reported that the speed of 8 – 9 knots when sailing was reduced by 6 – 7 knots when towing a beam trawl (Holdsworth 1874, 1883). Given the typical towing speed with the tide of 2 knot, a change in vessel speed of 6.5 knot would correspond to a gear resistance of 3.4 kN when towing at a speed of 1 knot above the tide. Gear resistance would be 2.8 kN when towing at 0.5 knot above the tide, and 4.3 kN when towing 1.5 knot above the tide. The estimate of the mechanic approach is about 40% lower than the gear resistance inferred from the semi quantitative information of Holdsworth.

Table 1. Dimensions of the vessel and gear components of a reference smack representative for the 1860s (Holdsworth, 1874). Parameter values are based on the dimensionless numbers and representative size of the vessel and beam trawl.

| Gear component | | Value | unit |
|---|---|---|---|
| Vessel | Length overall, LOA | 20 | m |
| | Length water line, $L$ | 18.2 | m |
| | Beam water line, $B$ | 5.1 | m |
| | Draught (mean), $T$ | 2.0 | m |
| | Displacement, $\nabla$ | 56 | $m^3$ |
| | Wetted surface, $A$ | 86 | $m^2$ |
| | Projected wetted surface in towing direction, $A_f$ | 38 | $m^2$ |
| | Longitudinal centre of buoyancy, *lcb* | 9.2 | m |
| | Midship coefficient, $CM$ | 0.653 | |
| | Block coefficient, $CB$ | 0.321 | |
| | Waterline coefficient, $CW$ | 0.701 | |
| | Prismatic coefficient, $CP$ | 0.473 | |
| | Friction coefficient, $C_f$ | 0.0053 | |
| Sails | Surface area | 205 | $m^2$ |
| Beam trawl | Length | 12 | m |
| | Diameter | 0.19 | m |
| | Weight | 1050 | $kg.m^{-3}$ |
| Shoes (2x) | Weight | 51.4 | kg |
| | Height | 1.05 | m |
| | Breath | 0.74 | m |
| | Width | 0.09 | m |
| | Frontal surface | 0.09 | $m^{-2}$ |
| | Footprint surface | 0.065 | $m^{-2}$ |
| Foot rope | Length | 20.0 | m |
| | Diameter | 0.15 | m |
| | Weight | 15.4 | $kg.m^{-1}$ |

| | | | |
|---|---|---|---|
| Head rope | Diameter | 0.05 | m |
| Netting | Twine surface area | 21.7 | $m^2$ |

Table 2. Parameter values used to estimate the hydrodynamic and geotechnical drag of the beam trawl gear components.

| Properties | | | | |
|---|---|---|---|---|
| Sea water | Specific weight | $\rho_{water}$ | 1021 | $kg.m^{-3}$ |
| Soil | Specific weight | $\rho_{soil}$ | 1700 | $kg.m^{-3}$ |
| Beam trawl | Beam (ash) | Specific weight | 1050 | $kg.m^{-3}$ |
| | Shoe (iron) | Specific weight | 7860 | $kg.m^{-3}$ |
| | Shoe | $C_d$ | 1.05 | |
| | Beam | $C_d$ | 1.1 | |
| | Warp | $C_d$ | 1.1 | |
| | Ground rope | $C_d$ | 1.1 | |
| | Ground rope | Weight per meter[2] | 21 | $kg.m^{-1}$ |
| | Ground rope | Specific weight | 1500 | $kg.m^{-3}$ |
| Beam trawl | | K1 | 0.47 | |
| | | K2 | 2.6 | |
| | | K3 | 0 | |

[2] Calculated with data from https://www.engineeringtoolbox.com/manila-rope-strength-d_1512.html. Downloaded 16 March 2025.

#### Acknowledgments

We gratefully acknowledge the help and critical feedback from drs Wick Hillege (Dykstra-NA), J. Hernandez Montfort, M. Verhulst (MARIN), G. Jacobi (Delft-University), prof dr Ana Ivanovich, dr F. O'Neill and prof dr J. van Leeuwen (Wageningen University).

Holdsworth EWH (1874) Deep-sea fishing and fishing boats. Edwards Stanford, London, England

Holdsworth EWH (1883) The sea fisheries of Great Britain and Ireland: an account of the practical working of the various fisheries around the British Islands. E. Stanford, London

Holtrop J, Mennen GGJ (1982) An approximate power prediction method. Int Shipbuild Prog 29:166–170

Ivanović A, Neilson RD, O'Neill FG (2011) Modelling the physical impact of trawl components on the seabed and comparison with sea trials. Ocean Eng 38:925–933

Larsson LC, Eliasson RE, Orych M (2022) Principles of yacht design. Adlard Coles, Bloomsbury Pubishing Plc., london, England

O'Neill FG, Ivanović A (2016) The physical impact of towed demersal fishing gears on soft sediments. ICES Journal of Marine Science 73:i5–i14

O'Neill FG, Summerbell K, Ivanović A (2018) The contact drag of towed demersal fishing gear components. J Mar Syst 177:39–52

O'Neill FG, Summerbell KJ (2016) The hydrodynamic drag and the mobilisation of sediment into the water column of towed fishing gear components. J Mar Syst 164:76–84

Paschen M, Richter U, Kopnick W (2000) Trawl penetration in the seabed (TRAPESE). Final Report EC-Study Contract No 96-006 University of Rostock, Rostock, Germany:150

Reid AJ (1977) A net drag formula for pelagic nets. Scottish Fisheries Research Report. Number 7.

Rijnsdorp AD, Depestele J, Molenaar P, Eigaard OR, Ivanovič A, O'Neill FG (2021) Sediment mobilisation by bottom trawls: a model approach applied to the Dutch North Sea beam trawl fishery ICES Journal of Marine Science 78:1574–1586

Slooff JW (2015) The aero- and hydromechanics of keel yachts. Springer International Publishing
